# Investigating the Genetic Architecture of Non-Cognitive Skills Using GWAS-by-Subtraction

**DOI:** 10.1101/2020.01.14.905794

**Authors:** Perline A. Demange, Margherita Malanchini, Travis T. Mallard, Pietro Biroli, Simon R. Cox, Andrew D. Grotzinger, Elliot M. Tucker-Drob, Abdel Abdellaoui, Louise Arseneault, Avshalom Caspi, David Corcoran, Benjamin Domingue, Colter Mitchell, Elsje van Bergen, Dorret I. Boomsma, Kathleen M. Harris, Hill F. Ip, Terrie E. Moffitt, Richie Poulton, Joseph Prinz, Karen Sugden, Jasmin Wertz, Benjamin Williams, Eveline L. de Zeeuw, Daniel W. Belsky, K. Paige Harden, Michel G. Nivard

**Affiliations:** Department of Biological Psychology, Vrije Universiteit Amsterdam, The Netherlands; Amsterdam Public Health Research Institute, Amsterdam University Medical Centers, Amsterdam, The Netherlands; Research Institute LEARN!,Vrije Universiteit Amsterdam, The Netherlands; Department of Biological and Experimental Psychology, Queen Mary University of London, UK; Social, Genetic and Developmental Psychiatric Centre, Institute of Psychiatry, King’s College London, UK; Department of Psychology, University of Texas at Austin, USA; Department of Economics, University of Zurich, Switzerland; Centre for Cognitive Ageing and Cognitive Epidemiology, University of Edinburgh, UK; Population Research Center, University of Texas at Austin, USA; Department of Psychiatry, Academic Medical Center, University of Amsterdam, Amsterdam, the Netherlands; Department of Psychology & Neuroscience, Duke University, Durham, NC, USA; Department of Psychiatry and Behavioral Sciences, Duke University School of Medicine, Durham, NC, USA; Center for Genomic and Computational Biology, Duke University, Durham, NC, USA; Stanford Graduate School of Education, Stanford University, Palo Alto, CA, USA; Institute for Social Research, University of Michigan, Ann Arbor, USA; Carolina Population Center and Department of Sociology, University of North Carolina at Chapel Hill, Chapel Hill, NC, USA; Department of Psychology and Dunedin Multidisciplinary Health and Development Research Unit, University of Otago, Otago, NZ; Department of Epidemiology, Columbia University Mailman School of Public Health, New York, NY, USA; Robert N. Butler Columbia Aging Center, Columbia University, New York, NY, USA

## Abstract

Educational attainment (EA) is influenced by cognitive abilities and by other characteristics and traits. However little is known about the genetic architecture of these “non-cognitive” contributions to EA. Here, we use Genomic Structural Equation Modelling and results of prior genome-wide association studies (GWASs) of EA (N = 1,131,881) and cognitive test performance (N = 257,841) to estimate SNP associations with variation in EA that is independent of cognitive ability. We identified 157 genome-wide significant loci and a polygenic architecture accounting for 57% of genetic variance in EA. Phenotypic and biological annotation revealed that (1) both cognitive and non-cognitive contributions to EA were genetically correlated with socioeconomic success and longevity; and (2) non-cognitive contributions to EA were related to personality, decision making, risk-behavior, and increased risk for psychiatric disorders; (3) non-cognitive and cognitive contributions to EA were enriched in the same tissues and cell types, but (4) showed different associations with gray-matter neuroimaging phenotypes.

---

Success in school – and in life – depends on skills beyond cognitive ability^1–4^. Randomized trials of early-life education interventions find substantial benefits to educational outcomes, employment, and adult health, even though the interventions have no lasting effects on children’s cognitive functions^5, 6^. These results have captured the attention of educators and policy makers, motivating growing interest in so-called “non-cognitive skills”^7–9^. Among non-cognitive skills suspected to be important for educational success are motivation, curiosity, persistence, and self-control^1,10–13^. However, questions have been raised about the substance of these skills and the magnitudes of their impacts on life outcomes^14^.

Twin studies find evidence that non-cognitive skills are heritable^3,15–18^. Genetic analysis could help clarify the contribution of these skills to educational attainment and elucidate their connections with other traits. A challenge to genetic research is a lack of consistent and reliable measurements of non-cognitive skills in existing genetic datasets^19^.

To overcome this challenge, we borrowed the strategy used in the original analysis of non-cognitive skills within the discipline of economics^20,21^: We operationalized non-cognitive skills as a latent variable that reflects the joint influence of all traits *other* than cognitive ability that contribute to educational attainment. We applied Genomic Structural Equation Modeling (Genomic-SEM)^22^ to existing GWASs of EA and cognitive performance (CP)^23^ in order to conduct a GWAS-by-subtraction. This approach allows us to estimate genetic associations with a non-cognitive skills phenotype that was never directly measured.

To evaluate results of the GWAS-by-subtraction of non-cognitive skills, we conducted phenotypic and biological annotation analysis. We used genetic correlation and polygenic score analysis to test genetic associations between non-cognitive skills and an array of socioeconomic and health outcomes, and relevant individual differences suggested by literature from different research fields. We also performed biological annotation analyses in order to identify cell types, tissues, and neurobiological structures that differentially relate to cognitive and non-cognitive skills.

## Results

### GWAS-by-Subtraction Identifies Genetic Associations with Non-Cognitive Variance in Educational Attainment

The term “non-cognitive skills” was originally coined by economists studying individuals who were equivalent in cognitive ability, but who differed in educational attainment.^21^ Our analysis of non-cognitive skills was designed to mirror this original approach: We focused on genetic variation in educational outcomes not explained by genetic variation in cognitive ability. Specifically, we applied Genomic Structural Equation Modeling (Genomic-SEM)^22^ to summary statistics from GWASs of educational attainment (EA)^23^ and CP^23^ (**Figure 1**). Both EA and CP were regressed on a latent factor, which captured genetic variance in CP (hereafter “*Cog*”). EA was further regressed on a second latent factor capturing genetic variance in EA independent of CP, hereafter “*NonCog*”. By construction, genetic variance in *NonCog* was independent of genetic variance in *Cog* (*r*g=0). In other words, the *NonCog* factor represents residual genetic variation in educational attainment that is not accounted for by the *Cog* factor. These two latent factors were then regressed on individual SNPs, yielding a GWAS of the latent constructs *Cog* and *NonCog*.

**Figure 1.**
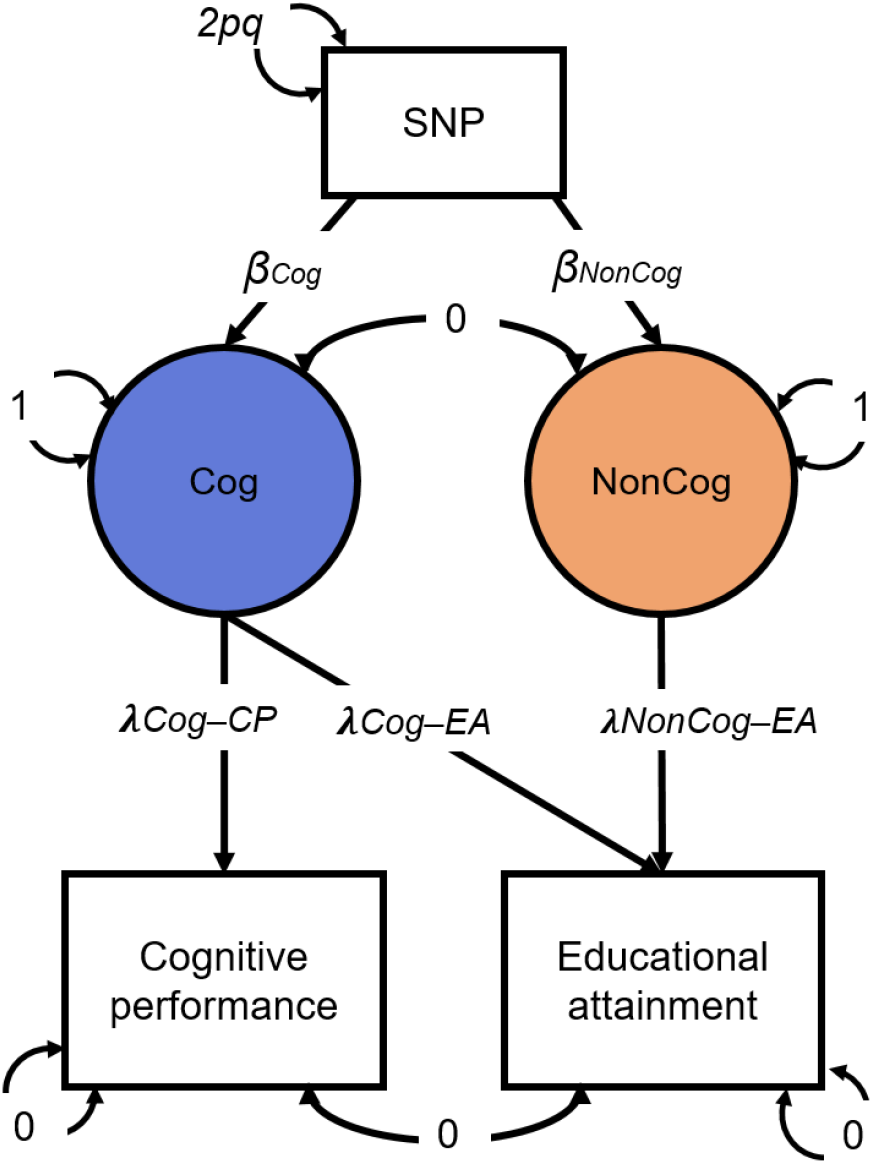
GWAS-by-subtraction Genomic-SEM model. Cholesky model as fitted in Genomic SEM, with path estimates for a single SNP included as illustration. SNP, Cognitive performance (CP) and Educational attainment (EA) are observed variables based on GWAS summary statistics. The genetic covariance between CP and EA is estimated based on GWAS summary statistics for CP and EA. The model is fitted to a 3×3 observed variance-covariance matrix (i.e. SNP, CP, EA). Cog and Non-Cog are latent (unobserved) variables. The covariances between CP and EA and between Cog and NonCog are fixed to 0. The variance of the SNP is fixed to the value of 2pq (p = reference allele frequency, q = alternative allele frequency, based on 1000 Genomes phase 3). The variances of CP and EA are fixed to 0, so that all variance is explained by the latent factors. The variances of the latent factors are fixed to 1. The observed variables CP and EA were regressed on the latent variables resulting in the estimates for the path loadings: λCog-CP=.4465; λCog-EA=.2305; λNonCog-EA=.2432. The latent variables were then regressed on each SNP that met QC criteria.

The *NonCog* latent factor accounted for 57% of genetic variance in EA. LD Score regression analysis estimated the *NonCog* SNP-heritability as *h*^2^_*NonCog*_=.0637 (*SE*=.0021). After Bonferroni correction, GWAS of *NonCog* identified 157 independent genome-wide significant lead SNPs (**Figure 2**) (independent SNPs defined as outside a 250Kb window, or within a 250Kb window and *r*^2^ < 0.1). As SNP associations on CP are entirely mediated by the *Cog* latent factor, results from the GWAS of *Cog* parallel the original GWAS of CP reported by Lee et al. (2018)^23^ and are reported in **Supplementary Note 1**.

**Figure 2.**
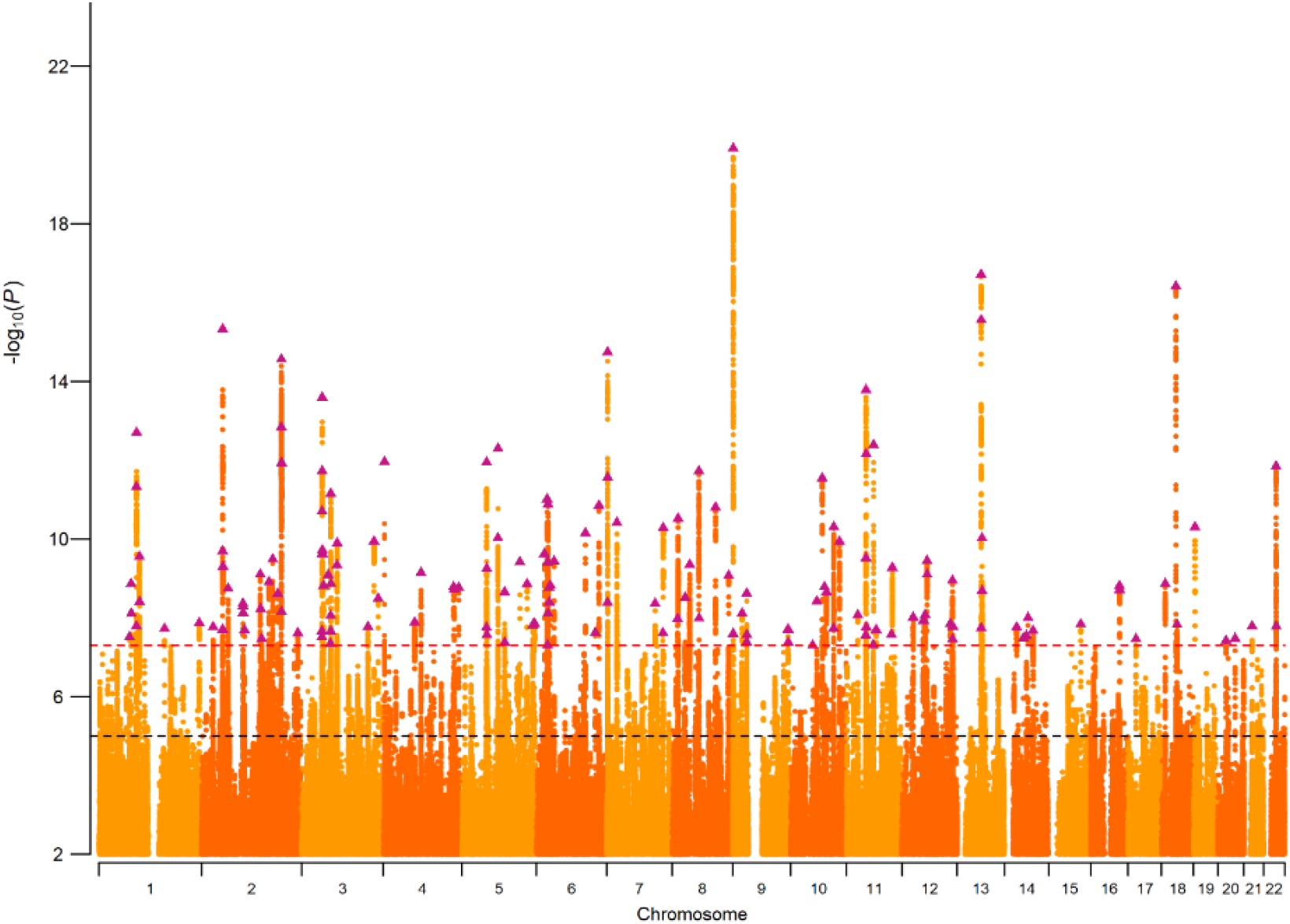
Manhattan plot of SNP associations with NonCog. Plot of the −log_10_(p-value) associated with the Wald test of β_NonCog_ for all SNPs, ordered by chromosome and base position. Purple triangles indicate genome-wide significant (p < 5e10^−8^) and independent (within a 250Kb window and r^2^ < .1) associations.

### Phenotypic Annotation I: Validating the *NonCog* Factor

To establish if the Genomic-SEM GWAS-by-subtraction succeeded in isolating genetic variance in education that was independent of cognitive function, we investigated whether *NonCog* genetically correlated with measures related to educational attainment and cognitive function and compared genetic correlations with *Cog*. We also confirm these results by conducting polygenic score (PGS) meta-analysis in 6 independent cohorts from the Netherlands (Netherlands Twin Register^24^ [NTR]), the U.S. (Texas Twin Project^25^; National Longitudinal Study of Adolescent to Adult Health^26^ [AddHealth], Wisconsin Longitudinal Study^27^ [WLS]), New Zealand (Dunedin Longitudinal Study^28^), and the United Kingdom (E-Risk^29^). Results reported in this preprint omit polygenic score analysis of the Dunedin and E-Risk cohorts. Effect-sizes for *r_g_* and PGS analysis are reported in **Figure 3 and Supplementary Figure 2 and Supplementary Tables 4, 5 and 9**.

**Figure 3.**
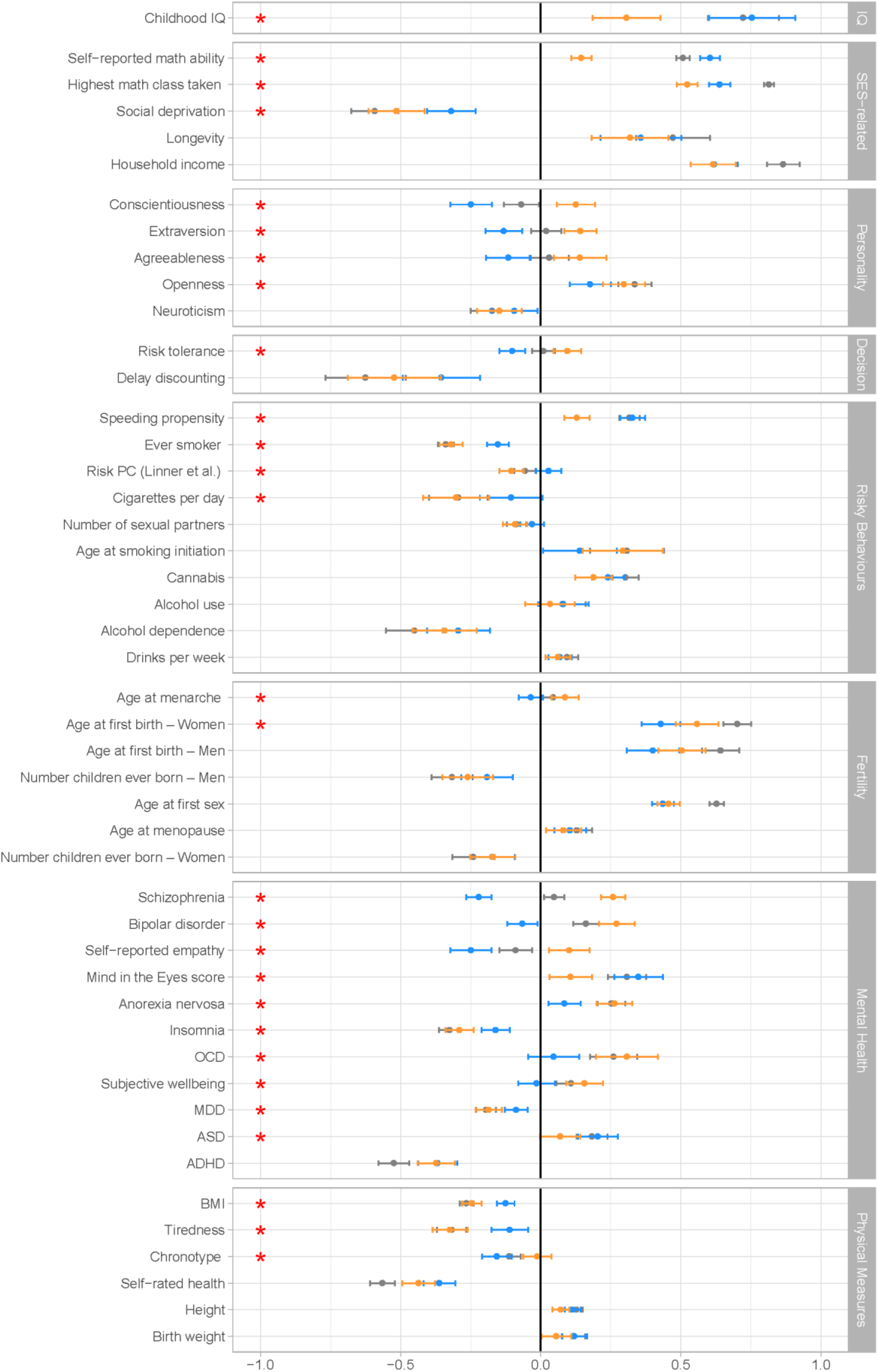
Estimates of genetic correlations with NonCog, Cog and Educational Attainment. Genetic correlations between NonCog, Cog, and EA and other traits of interest, as estimated with Genomic SEM. Correlations with NonCog are in orange; with Cog in blue; with EA in gray. Error bars represent 95% CIs. The red stars represents a significant (FDR corrected p-value < 0.05) difference in the magnitude of the correlation with Cog versus NonCog. The difference test is based on a chi-squared test associated with a comparison between a model constraining these two correlations to be identical, versus a model where the correlations are freely estimated.

**Figure 4.**
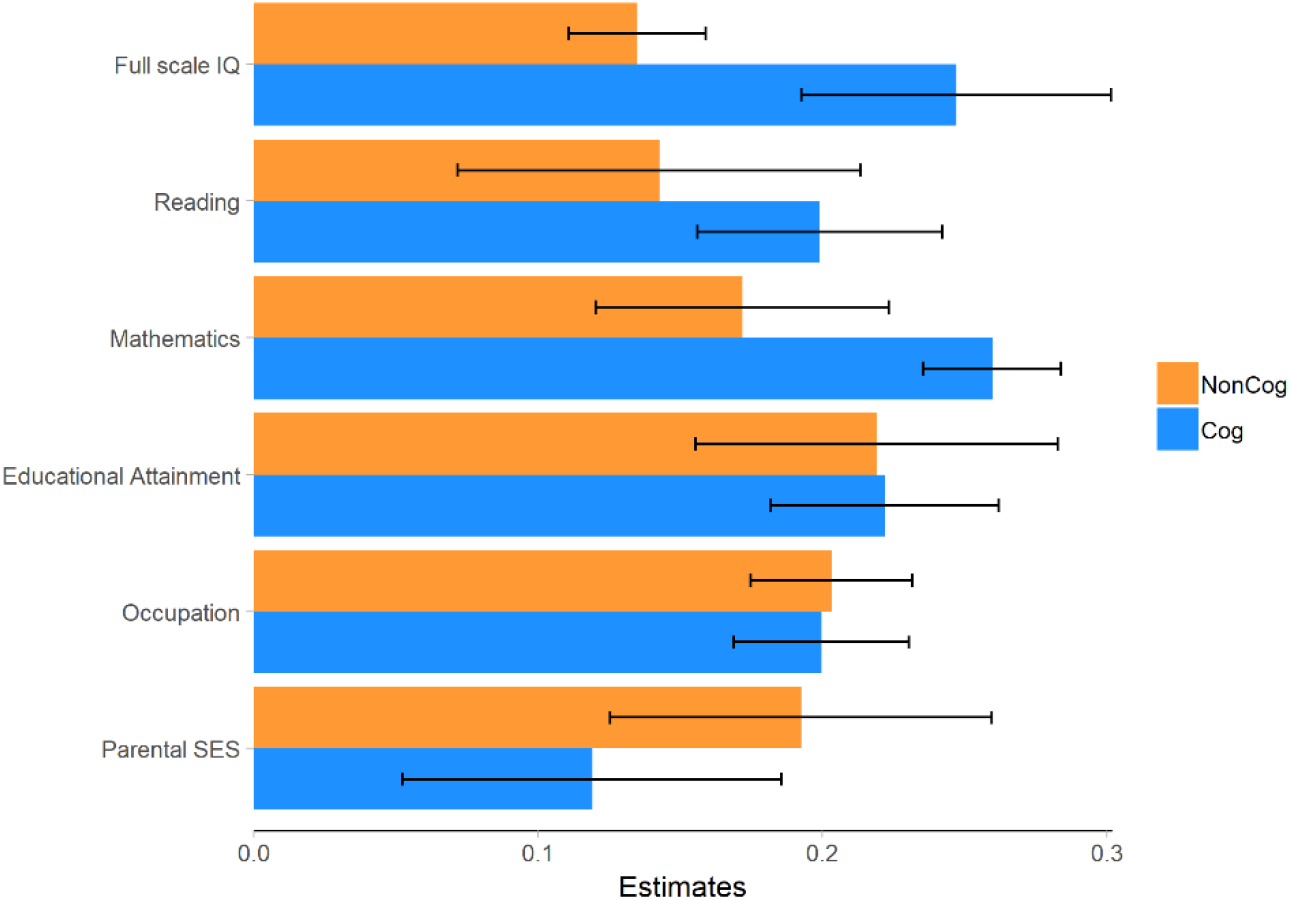
Polygenic prediction of IQ, achievement and socioeconomic measures. Meta-analytic estimates of the polygenic score associations with cognitive test performance, educational achievement and socioeconomic measures. Cog and NonCog PRS were entered simultaneously in multiple regression. 95% CI are represented by black bars. Cohorts, measures and sample size are detailed in Supplementary Tables 6-9. Traits were measured in different cohorts: Full scale IQ in WLS, Texas twins and NTR (N=2815); Reading and Mathematics Achievement in AddHealth, Texas Twins and NTR (N=9274-10474); Educational Attainment in AddHealth, WLS, and NTR (N=21365); Occupation in AddHealth (N=5527); Parental SES in Texas Twins (N=566).

*NonCog* genetics have weaker associations with cognitive functions as compared to *Cog* **genetics.** *NonCog* and *Cog* were both genetically correlated with childhood IQ^30^; however, the magnitude of *NonCog r*_g_ was less than half the *r*_g_ for *Cog* (*NonCog r*_g_=0.31 (*SE*=.06), *Cog r*_g_=0.75 (*SE*=.08), *p*_diff_fdr_<.0001). Of the total genetic correlation between childhood IQ and EA, 31% of the variance was explained by *NonCog* and 69% by *Cog*. In PGS analysis in the NTR and Texas Twin cohorts (*N*=2,815), effect-sizes for associations with IQ were smaller for *NonCog* as compared to *Cog* (*NonCog β*=.13 (*SE*=.01), *Cog β*=.25 (*SE*=.03); *p*_diff_<.0001; Dunedin and E-Risk analysis pending). Sensitivity analyses of tests measuring different dimensions of cognitive function are reported in **Supplementary Figure 2**. These results confirm that *NonCog* genetic associations with cognitive test performance, while greater than zero, are of smaller magnitude as compared to EA or *Cog* genetics.

### *NonCog* genetics have weaker associations with young people’s academic abilities/skills as compared to *Cog* genetics

We next tested if *NonCog* was genetically associated with academic abilities that contribute to educational attainment. *NonCog r*_g_ with self-reported math ability was positive and statistically different from zero, but smaller in magnitude as compared to *Cog* (*NonCog r*_g_=0.15 (*SE*=.02), *Cog r*_g_=0.61 (*SE*=.02), *p*_diff_fdr_<.0001). In Genomic-SEM analysis, *NonCog* explained 22% of the *r*_g_ between EA and math ability. In PGS analysis in the NTR, Texas-Twin, and AddHealth cohorts, *NonCog* and *Cog* polygenic scores were associated with reading and math skills, although effect-sizes were smaller for *NonCog* than for *Cog* (for reading: *NonCog β*=.14 (*SE*=.03), *Cog β*=.20 (*SE*=.02), *p*_diff_=.0032, *N*=9,274; for math: *NonCog β*=.17 (*SE*=.03), *Cog β*=.26 (*SE*=.01), *p*_diff_<.0001, *N*=10,474). These results suggested that *NonCog* skills are related to educational attainment in part through pathways other than the development of specific academic skills/abilities.

### *NonCog* genetics have similar associations with academic achievement as compared to *Cog* genetics

In contrast to difference between *NonCog* and *Cog* genetic correlations with self-reported math ability, genetic correlations were more similar for achievement in math education (self-report of most advanced math course taken: *NonCog r*_g_=0.52 (*SE*=.02), *Cog r*_g_=0.64 (*SE*=.02), *p*_diff_fdr_<.0001). In Genomic SEM analysis, NonCog accounted for 48% of the *r*_g_ between EA and math achievement.

Findings were parallel for analysis of educational attainment. To compute *r*_*g*_ among *NonCog*, *Cog*, and educational attainment, we re-ran the Genomic-SEM model using summary statistics that omitted the 23andMe sample from the EA GWAS. We then computed the *r*_g_ between *NonCog* (estimated without 23andMe) and EA in the 23andMe sample. *NonCog* was more strongly associated with EA than was *Cog* (*NonCog r*_*g*_ =.71 (*SE*=.02), *Cog r_g_*=.57 (*SE*=.02), *p*_diff_ < .0001). In PGS analysis based on the full Genomic-SEM model including 23andMe, effect-sizes for associations with educational attainment were similar for *NonCog* and *Cog* (AddHealth, WLS and NTR meta-analysis *NonCog β*=.22 (*SE*=.03), *Cog β*=.22 (*SE*=.02), *p*_diff_=.63, total *N*=21,365; Dunedin and E-Risk analysis pending).

### *NonCog* genetics have similar associations with socioeconomic attainment and longevity as compared to *Cog* genetics

The public-health significance of educational attainment is partly due to its relationship with long-term economic and health outcomes^31,32^. We therefore tested if *NonCog* was related to these long-term outcomes and if magnitudes of associations were similar to those for *Cog*.

#### Socioeconomic Attainment

In genetic correlation analysis, *NonCog* was as strongly – or more strongly – associated with socioeconomic attainment outcomes, as compared to *Cog* (for income^33^, *NonCog r*_g_=.62, (*SE*=.04), *Cog r*_g_=.62 (*SE*=.04), *p*_diff_fdr_=.95; for neighborhood deprivation^33^, *NonCog r*_g_=−.51 (*SE*=.05), *Cog r*_g_=−.32 (*SE*=.04), *p*_diff_fdr_=.001). *NonCog* explained 53% of the EA *r*_g_ with income and 65% of the EA *r*_g_ with neighborhood deprivation. In PGS analysis in the AddHealth cohort (*N*=5,527), *NonCog* and *Cog* PGS showed similar associations with occupational attainment (*NonCog β*=.20 (*SE*=.01), *Cog β*=.20 (*SE*=.02), *p*_diff_=.865; Dunedin analysis pending).

#### Longevity

We estimated *r*_*g*_ with longevity as proxied by parental lifespan^34^. Genetic correlations were similar for *NonCog* and *Cog* (*NonCog r*_g_=.32 (*SE*=.07); *Cog r*_g_=.36 (*SE*=.07); *p*_diff_fdr_=.71). In Genomic-SEM analysis, *NonCog* explained 50% of the *r*_g_ between EA and longevity.

In sum, validation analysis found *NonCog* genetics were less-related to cognitive- and academic-ability phenotypes as compared to *Cog* genetics, but showed comparable associations with academic-, economic- and health-attainment phenotypes. These findings are consistent with GWAS-by-subtraction analysis having identified genetic influences on non-cognitive skills important to achievement in school and beyond.

### Phenotypic Annotation II. Exploring Genetic Correlates of the *NonCog* Factor

Our next set of phenotypic annotation analyses investigated connections between the *NonCog* factor and phenotypes linked with non-cognitive skills within the disciplines of economics and psychology (**Figure 3**).

### *NonCog* genetics were associated with decision-making preferences

In economics, non-cognitive influences on achievement and health are often studied in relation to decision-making preferences^35–38^. *NonCog* was genetically correlated with higher levels of comfort with risk-taking^39^ (risk tolerance *r*_g_=.10 (*SE*=.03)) and willingness to forego immediate gratification in favor of a larger reward at a later time^40^ (delay discounting *r*_g_=−.52 (*SE*=.08)). In contrast, *Cog* was genetically correlated with generally more cautious decision-making characterized by lower levels of risk tolerance (*r*_g_=−.35 (*SE*=.07), *p*_diff_fdr_<.0001) and moderate delay discounting (*r*_g_=−.10 (*SE*=.02), *p*_diff_fdr_=.0852).

### *NonCog* genetics were associated with less risky health behavior and delayed fertility

An alternative approach to studying non-cognitive skills in economics and other social sciences is to infer individual differences in non-cognitive skills from patterns of risk behavior. In genetic correlation analysis of obesity^41^, substance use^39,42–45^, and sexual behaviours and early fertility^39,46,47^, *NonCog* was consistently genetically correlated with lower levels of risk (*r*_g_ range .2-.5), with the exception that the *r*_g_ with alcohol use was not different from zero and *r*_g_ with cannabis use was positive. Genetic correlations for *Cog* were generally in the same direction but of smaller magnitude.

### *NonCog* genetics were associated with a broad spectrum of personality characteristics linked with social and professional competency

In psychology, non-cognitive influences on achievement are conceptualized as personality traits, *i.e.* patterns of stable individual differences in emotion and behavior. The model of personality that has received the most attention in genetics is a five-factor model referred to as the Big-5. Genetic correlation analysis of the Big-5 personality traits^48–50^ revealed *NonCog* genetics were most strongly associated with Openness to Experience (being curious and eager to learn; *r*_g_=.30 (*SE*=.04)) and were further associated with a pattern of personality characteristic of changes that occur as people mature in adulthood^51^. Specifically, *NonCog* showed a positive *r*_g_ with Conscientiousness (being industrious and orderly; *r*_g_=.13 (*SE*=.03)), Extraversion (being enthusiastic and assertive; *r*_g_=.14 (*SE*=.03)), and Agreeableness (being polite and compassionate; *r*_g_=.14 (*SE*=.05)), and negative *r*_g_ with Neuroticism (being emotionally volatile; *r*_g_=−.15 (*SE*=.04)). Genetic correlations of *Cog* with Openness to Experience and Neuroticism were similar to those for *NonCog* (*p*_diff_fdr-Openness_=.0414, *p*_diff_fdr-Neuroticism_=.4821). In contrast, genetic correlations of *Cog* with Conscientiousness, Extraversion, and Agreeableness were in the opposite direction (*r*_g_=−.12 to −.25, *p*_diff_fdr_<.0005).

We conducted PGS analysis of Big-5 personality in the NTR, Texas Twin, AddHealth, and WLS cohorts (*N* = 21,203 - 21,290 across personality traits) (**Supplementary Figure 3**). *NonCog* PGS associations with personality traits paralleled genetic correlations, but were smaller in magnitude and were statistically different from zero at the alpha=0.05 threshold only for Openness (meta-analytic *β*=.13 (*SE*=.02)) and Agreeableness (meta-analytic *β*=.04 (*SE*=.02)). Also parallel to genetic correlation analysis, the *Cog* PGS associations with openness and neuroticism were in the same direction but smaller in magnitude as compared to *NonCog* associations, and were in the opposite direction for conscientiousness, extraversion, and agreeableness, although only associations with openness, conscientiousness, and neuroticism were statistically different from zero at the alpha=0.05 level (meta-analytic *β*_Neuroticism_=−.05, *p*_diff_=<.0001; *β*_Openness_=.08, *p*_diff_=.152; *β*_Conscientiousness_=−.03, *p*_diff_=.001).

### *NonCog* genetics were associated with higher risk for multiple psychiatric disorders

In clinical psychology and psychiatry, research is focused on mental disorders. Mental disorders are generally associated with phenotypic impairments in academic achievement and social role functioning,^52,53^ but positive genetic correlations with educational attainment and creativity have been reported for some disorders^54,55^. We therefore tested *NonCog r*_*g*_ with psychiatric disorders based on published case-control GWAS^56–62^. *NonCog* was associated with *higher* risk for multiple clinically-defined disorders including anorexia nervosa (*r*_g_=.26 (*SE*=.04)), obsessive-compulsive disorder (*r*_g_=.31 (*SE*=.06)), bipolar disorder (*r*_g_=.27 (*SE*=.03)), and schizophrenia (*r*_g_=.26 (*SE*=.02)). Genetic correlations between *Cog* and psychiatric disorders were either much smaller in magnitude (anorexia nervosa *r*_g_=.08 (*SE*=.03), *p*_diff_fdr_<.001; obsessive-compulsive disorder *r*_g_=.05 (*SE*=.05), *p*_diff_fdr_<.01) or in the opposite direction (bipolar disorder *r*_g_=−.07 (*SE*=.03), *p*_diff_fdr_<.001; schizophrenia *r*_g_=−.22 (*SE*=.02), *p*_diff_fdr_<.001). Both *NonCog* showed negative genetic correlations with attention-deficit/hyperactivity disorder (*NonCog r*_g_=−.37 (*SE*=.03), *Cog r*_g_=−.37 (*SE*=.04), *p*_diff_fdr_=.95).

In sum *NonCog* genetics were associated with phenotypes from economics and psychology thought to mediate non-cognitive influences on educational success. These associations contrasted with associations for *Cog* genetics, supporting distinct pathways of influence on achievement in school and later in life. Opposing patterns of association were also observed for psychiatric disorders, suggesting that the unexpected positive genetic correlation between educational attainment and mental health problems uncovered in previous studies arises from non-cognitive genetic influences on educational attainment.

### Biological Annotation Analysis Reveal Shared and Specific Neurobiological Correlates *NonCog* and *Cog* genetics were enriched in similar tissues and cells

We tested whether common variants in genes specifically expressed in 53 GTEx tissues^63^ or in 152 tissues captured in a previous aggregation of RNA-seq studies^64,65^ were enriched in their effects on *Cog* or *NonCog*. Genes predominantly expressed in the brain rather than peripheral tissues were enriched in both *NonCog* and *Cog* (**Supplementary Table 10**).

To examine expression patterns at a more granular level of analysis, we used MAGMA^66^ and stratified LD score regression^67^ to test enrichment of common variants in 265 brain cell-type-specific gene-sets^68^. In MAGMA analysis, common variants in 95 of 265 gene-sets were enriched for association with *NonCog*. The enriched cell-types were predominantly neurons (97%), with enrichment most pronounced for telencephalon-projecting neurons, di- and mesencephalon neurons, and to a lesser extent, telencephalon interneurons (**Supplementary Figure 4** and **Table 12**). As measured by correlation between *Z*-statistics, enrichment for *Cog* was similar to *NonCog* (*r*=.85) and there were no differences in cell-type-specific enrichment, suggesting little differentiation between cognitive ability and non-cognitive traits at the level of cell-type (**Supplementary Figure 5**). Stratified LDSC results were similar to results from MAGMA (**Supplementary Note 2, Supplementary Figure 6** and **Table 13**). While the same gene-sets, based on scRNA-seq expression in neuronal cell-types, are enriched for *NonCog* and *Cog*, gene-level analysis^69^ (**Supplementary Note 3**) confirms the specific genes driving this enrichment do not necessarily affect the two traits in the same direction.

### *NonCog* and *Cog* genetics show diverging associations with total and regional brain volumes

EA is genetically correlated with greater total brain volume^70,71^. We therefore compared the *r*_g_ of *NonCog* and *Cog* with total brain volume and with 100 regional brain volumes (99 gray matter volumes and white matter volume) controlling for total brain volume (**Supplementary Table 15**)^72^. For total brain volume, genetic correlation was stronger for *Cog* as compared to *NonCog* (*Cog r*_g_=.22 (*SE*=.04), *NonCog r*_g_=.07 (*SE*=.03), *p*_diff_=.005). Total gray matter volume, controlling for total brain volume, was not associated with either *NonCog* or *Cog* (*NonCog*: *r*_g_=.07 (*SE*=.04); *Cog*: *r*_g_=.06 (*SE*=.04)). For total white matter volume, conditional on total brain volume, genetic correlation was negative and stronger for *NonCog* as compared to *Cog* (*NonCog r*_g_= −.12 (*SE*=.04), *Cog* (*r*_g_=−.01 (*SE*=.04), *p*_diff_=.04).

*NonCog* was not associated with any of the regional gray-matter volumes after FDR correction. In contrast, *Cog* was significantly associated with regional gray-matter volumes for the bilateral fusiform, insula and posterior cingulate (*r*_g_ range .11-.17), as well as left superior temporal (*r*_g_=.11 (*SE*=.04)), left pericalcarine (*r*_g_=−.16 (*SE*=.05)) and right superior parietal volumes (*r*_g_=−.22 (*SE*=.06)) (**Figure 5**).

**Figure 5.**
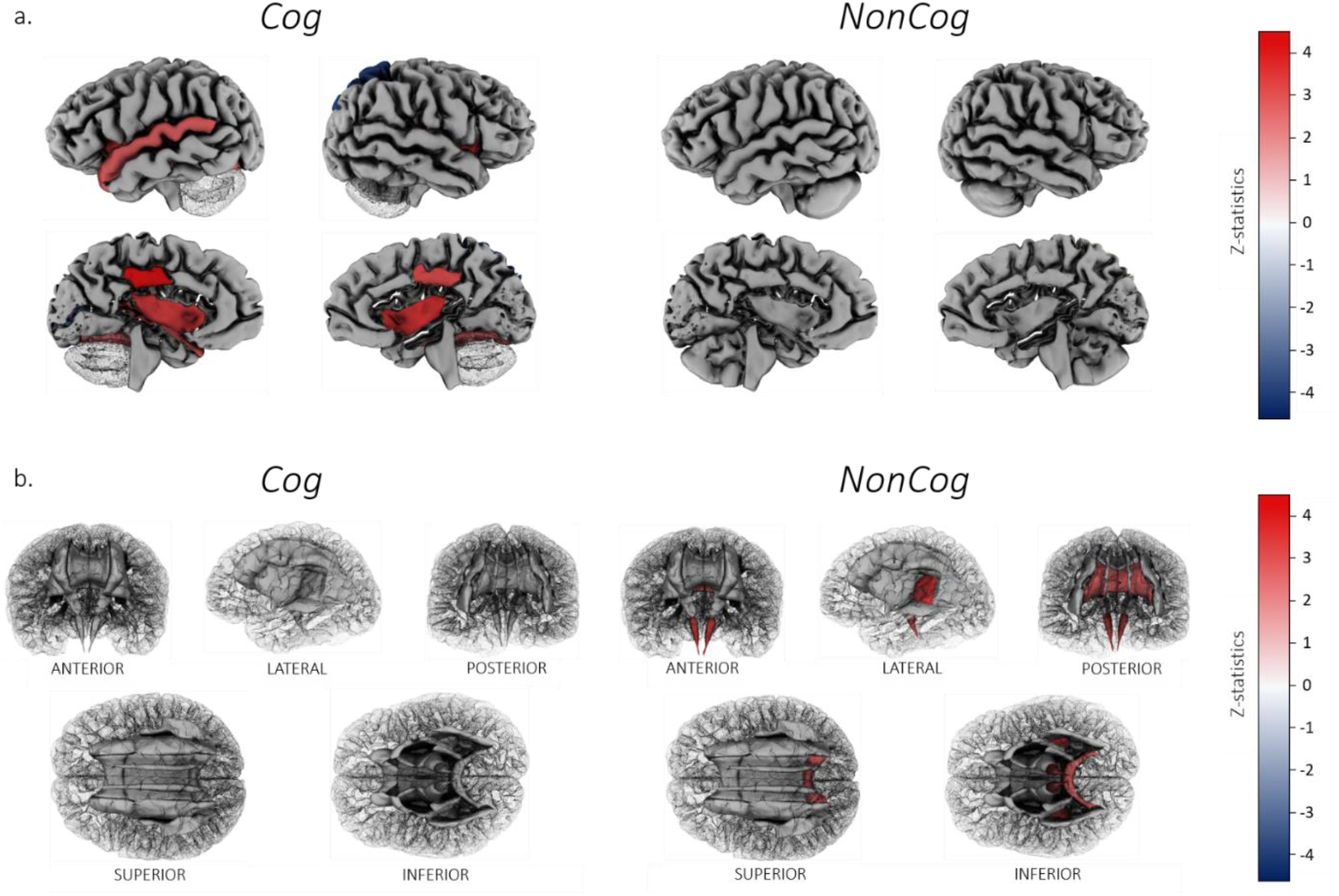
Genetic correlations with regional gray matter volumes and white matter tracts. a. Cortical patterning of FDR-corrected significant genetic correlations with regional gray matter volumes for Cog versus NonCog, after correction for total brain volume. Regions of interest are plotted according to the Desikan-Killiany-Tourville atlas, shown on a single manually-edited surface (Klein & Tourville, 2012; http://mindboggle.info). Cog showed significant associations with gray matter volume for the bilateral fusiform, insula and posterior cingulate, the left superior temporal and left pericalcarine and right superior parietal volumes. NonCog was not associated with any of the regional brain volumes. b. White matter tract patterning of FDR-corrected significant genetic correlations with regional mode of anisotropy (MO) for Cog versus NonCog. White matter tract probability maps are plotted according to the Johns Hopkins University DTI atlas (https://neurovault.org/). Cog was not associated with regional MO. NonCog showed significant associations with MO in the corticospinal tract, the retrolenticular limb of the internal capsule and the splenium of the corpus callosum.

### *NonCog* and *Cog* genetics were weakly associated with white matter microstructure

We tested genetic correlation of *NonCog* and *Cog* with white matter tract integrity as measured using diffusion tensor imaging (DTI)^73^. Analyses included 5 DTI parameters in each of 22 white matter tracts (**Supplementary Table 16**): fractional anisotropy (FA), mean diffusivity (MD), axial diffusivity (AD), radial diffusivity (RD), and mode of anisotropy (MO).

We first analyzed tract-wide association of *NonCog* and *Cog* with the five DTI parameters. *Cog* was nominally associated with global white matter microstructure for two DTI parameters – average AD (*r*_g_=.09 (*SE*=.04); greater diffusion of water along the principal axis of diffusion) and average MO (*r*_g_=.11 (*SE*=.04); more tubular, as opposed to planar, water diffusion). Only average MO survived FDR correction (*q*=.014). These genetic correlations did not differ from genetic correlations with the *NonCog* factor (χ^2^ *p* >.324).

Next, we analyzed tract-specific genetic correlations for each of the 5 DTI parameters. *NonCog* was positively associated with MO in the corticospinal tract (*r*_g_=.14 (*SE*=.05)), retrolenticular limb of the internal capsule (*r*_g_=.12 (*SE*=.04)) and splenium of the corpus callosum (*r*_g_=.10 (*SE*=.04); **Figure 5**), whereas the *Cog* factor was not associated with any specific tracts. However, none of the FDR-significant associations for *NonCog* were statistically different from associations for *Cog* (*p*_diff_fdr_=.89-.99), possibly reflecting a lack of power to detect differences in small effects.

## Discussion

GWAS of non-cognitive influences on educational attainment (EA) identified 157 independent loci and polygenic architecture accounting for more than half the genetic variance in EA. In genetic correlation and PGS analysis, these non-cognitive (*NonCog*) genetics showed similar magnitude of associations with EA, economic attainment and longevity to genetics associated with cognitive influences on EA (*Cog*). As expected, *NonCog* genetics had much weaker associations with cognition phenotypes as compared to *Cog* genetics. These results contribute new GWAS evidence in support of the hypothesis that heritable non-cognitive skills influence educational attainment and downstream life-course economic and health outcomes.

Phenotypic and biological annotation analyses shed light on the substance of heritable non-cognitive skills influencing education. Economists hypothesize that preferences that guide decision-making in the face of risk and delayed rewards represent non-cognitive influences on educational attainment. Consistent with this hypothesis, *NonCog* genetics were associated with higher risk tolerance and lower time discounting. These decision-making preferences are associated with financial wealth, whereas opposite biases are hypothesized to contribute to a feedback loop perpetuating poverty^74^. Consistent with results from analysis of decision-making preferences, *NonCog* genetics were also associated with healthier behavior and later fertility.

Psychologists hypothesize that the Big Five personality characteristics of conscientiousness and openness are the two “pillars of educational success”^2,3,75^. Our results provide some support for this hypothesis, with the strongest genetic correlation evident for openness. But they also show that non-cognitive skills encompass the full range of personality traits, including agreeableness, extraversion, and the absence of neuroticism. This pattern mirrors the pattern of personality change that occurs as young people mature into adulthood^51^. Thus, non-cognitive skills share genetic etiology with what might be termed as “mature personality”. The absolute magnitudes of genetic correlations between *NonCog* and individual personality traits are modest. This result suggests that the personality traits described by psychologists capture some, but not all genetic influence on non-cognitive skills.

Although the general pattern of findings in our phenotypic annotation analysis indicated non-cognitive skills were genetically related to socially desirable characteristics and behaviors, there was an important exception. Genetic correlation analysis of psychiatric disorder GWAS revealed positive associations of *NonCog* genetics with schizophrenia, bipolar disorder, anorexia nervosa, and obsessive-compulsive disorder. Previously, these psychiatric disorders have been shown to have a positive *r*_g_ with EA, a result that has been characterized as paradoxical given the impairments in educational and occupational functioning typical of serious mental illness. Our results clarify that these associations are driven by non-cognitive factors associated with success in education. These results align with the theory that clinically-defined psychiatric disorders represent extreme manifestations of dimensional psychological traits, which might be associated with adaptive functioning within the normal range^76–78^.

Our analysis found little support for the hypothesis that physical attributes, such as attractive appearance, might be associated with academic success because of social biases.^79^ There are not yet well-powered GWAS of physical attractiveness. However, tissue enrichments for *NonCog* genetics were found only in the brain and not in any peripheral tissues and genetic correlation with stature, a generally socially desirable physical attribute, were much smaller than for behavioral and psychological phenotypes.

Finally, biological annotation analyses suggest similarities in the cellular mediators of *NonCog* and *Cog* influences on educational attainment. In tissue-enrichment analysis, GWAS results for both *Cog* and *NonCog* were enriched for gene-sets predominantly expressed in the brain. In gene-set enrichment analysis, there were no statistically significant differences between *NonCog* and *Cog.* Thus, while the effects of genetic variation on *NonCog* and *Cog* are uncorrelated, the variation resides in the same (functional) regions of the genome that play a role in specific types of neurons. We found some evidence of differences between *NonCog* and *Cog* in associations with brain structure: *NonCog* was more strongly associated with white matter microstructure as compared to *Cog*, whereas *Cog* was more strongly associated with gray matter volumes as compared to *NonCog*. Moderate sample sizes in neuroimaging GWAS mean these results must be treated as preliminary, requiring replication with data from larger-scale GWAS of white-matter and gray-matter phenotypes. Results also illustrate how Genomic-SEM can be used to conduct GWAS of phenotypes not directly measured in large-scale databases, an application that might have broad utility beyond the genetics of educational attainment.

We acknowledge limitations. Genomic-SEM analysis to isolate non-cognitive genetic influences on educational attainment relies on a statistical model of a complex developmental process. Cognitive and non-cognitive skills develop in interaction with one another. For example, the dynamic mutualism hypothesis^80^ proposes that non-cognitive characteristics shape investments of time and effort, leading to differences in the pace of cognitive development^81,82^. In Genomic-SEM analysis, the *NonCog* factor is, by construction, uncorrelated with adult cognition. Thus, the statistical model is an imperfect representation of etiology. Nevertheless, statistical separation of *NonCog* from *Cog*, although artificial, allows us to test if heritable traits other than cognitive ability influence educational attainment and to explore what those traits may be. Our finding that *NonCog* genetics account for roughly half of all genetic variance in EA should motivate future longitudinal studies to collect repeated measures of cognitive and non-cognitive skills in order to study their reciprocal relationship across development^83,84^.

Our use of Genomic-SEM to perform GWAS-by-subtraction relied on published GWASs of adult cognitive performance and of educational attainment. Biases and limitations in these GWASs will also affect our results. For example, a large portion of data in the cognitive performance GWAS came from UK Biobank, which administered only a limited battery of cognitive tests. This limited battery could fail to capture genetic influences on some cognitive functions, resulting in incomplete separation of cognitive from non-cognitive genetics within the Genomic-SEM analysis. Genomic-SEM analysis of *NonCog* genetics using data from GWAS with more comprehensive cognitive testing is needed.

In the case of GWAS of educational attainment, the included samples were drawn mainly from Western Europe and the U.S., and participants completed their education in the late 20^th^ and early 21^st^ centuries. The phenotype of educational attainment reflects an interaction between an individual and the social system in which they are educated. Differences across social systems, including education policy, culture, and historical context, may result in different heritable traits having influence on educational attainment^85^. As a result, the GWAS results for educational attainment and the Genomic-SEM results for non-cognitive skills based on these results may not generalize beyond the times and places when and where GWAS samples were collected. Follow-up analysis in cohorts drawn from other contexts are needed to clarify how findings for *NonCog* genetics generalize.

Generalization of the *NonCog* factor is also limited by the homogeneity of ancestry in the educational attainment and cognitive performance GWASs. Analysis included only participants of European descent. Although this restricted sample is necessary given the lack of methods for integrating genome-scale genetic data across populations with different ancestries^86,87^, it raises a potential threat to external validity. Analysis of (*Non)Cog* in non-European populations should be a priority following either the conduct of GWAS in other ancestries or the refinement of methods to better integrate data across samples drawn from different ancestries.

Within the bounds of these limitations, our analysis provides a first view of the genetic architecture of non-cognitive skills influencing educational success. These skills are central to theories of human capital formation within the social and behavioral sciences and are increasingly the targets of social policy interventions. Our results establish that non-cognitive skills are central to the heritability of educational attainment and illuminate connections between genetic influences on these skills and social and behavioral science phenotypes.

## Methods

### Meta-analysis of educational attainment GWAS

We reproduced the Social Science Genetic Association Consortium (SSGAC) 2018 GWAS of educational attainment^23^ by meta-analyzing published summary statistics for *N*=766,345 (www.thessgac.org/data) with summary statistics obtained from 23andMe, Inc. (N=365,538). We included SNPs with sample-size > 500,000 and MAF > 0.005 in the 1000 Genomes reference set (10,101,243 SNPs). We did not apply genomic control, as standard errors of publicly available and 23andMe summary statistics were already corrected^23^. Meta-analysis was performed using METAL^88^.

### GWAS-by-subtraction

The objective of our GWAS-by-subtraction analysis was to estimate, for each SNP, the association with educational attainment that was independent of that SNP’s association with cognition (hereafter, the *NonCog* SNP effect). We used Genomic-SEM^22^ to analyze GWAS summary statistics for the educational attainment and cognitive performance phenotypes in the SSGAC’s 2018 GWAS (Lee et al. 2018^23^). The model regressed the educational-attainment and cognitive-performance summary statistics on two latent variables, *Cog* and *NonCog* (**Figure 1**). *Cog* and *NonCog* were then regressed on each SNP in the genome. This analysis allowed for two paths of association with educational attainment for each SNP. One path was fully mediated by *Cog*. The other path was independent of *Cog* and measured the non-cognitive SNP effect, *NonCog*. To identify independent lead hits with *p* <5e-8 (the customary p-value threshold to approximate an alpha value of 0.05 in GWAS), we pruned the results using a radius of 250 kb and an LD threshold of r^2^ <0.1 (**Supplementary Tables 1 and 2**).

### Genetic correlations

We use Genomic-SEM to compute genetic correlations of *Cog* and *NonCog* with other education-linked traits for which well-powered GWAS data were available (SNP-*h*^2^ z-score >2; **Supplementary Table 3**) and to test if genetic correlations with these traits differed between *Cog* and *NonCog*. Specifically, models tested the null hypothesis that trait genetic correlations with *Cog* and *NonCog* could be constrained to be equal using a chi-squared test with FDR adjustment to correct for multiple testing. The FDR adjustment was conducted across all genetic correlation analyses reported in the article excluding the analyses of brain volumes described below. Finally, we used Genomic-SEM analysis of genetic correlations to estimate the percentage of the genetic covariance between educational attainment and the target traits that was explained by *Cog* and *NonCog* using the model illustrated in **Supplementary Figure 8**.

### Polygenic score analysis

Polygenic score analyses were conducted in data drawn from six population-based cohorts from the Netherlands, the U.K., the U.S., and New Zealand: (1) the Netherlands Twin Register (NTR)^24,89^, (2) E-Risk^29^, (3) the Texas Twin Project^25^, (4) the National Longitudinal Study of Adolescent to Adult Health (AddHealth)^26,90^, dbGaP accession phs001367.v1.p1; (5) Wisconsin Longitudinal Study on Aging (WLS)^27^, dbGaP accession phs001157.v1.p1; and (6) the Dunedin Multidisciplinary Health and Development Study^28^. (At the time this preprint was posted, Dunedin and E-Risk analyses were not yet complete and data from these studies is not included in the reported analysis.) **Supplementary Tables 6 and 7** describe cohort-specific metrics. Polygenic scores were computed based on weights derived using the LD-pred^91^ software with an infinitesimal prior and the 1000 Genomes phase 3 sample as a reference for the LD structure. LD-pred weights were computed in a shared pipeline to ensure comparability between cohorts. Each outcome (*e.g.*, IQ score) was regressed on the *Cog* and *NonCog* polygenic scores and a set of control variables (sex, 10 principal components derived from the genetic data and, for cohorts in which these quantities varied, genotyping chip and age). In cohorts containing related individuals, non-independence of observations from relatives were accounted for using mixed linear models (MLM), generalized estimation equations (GEE), or by clustering of standard errors at the family level. We used a random effects meta-analysis to aggregate the results across the cohorts. This analysis allows a cohort-specific random intercept. Individual cohort results are in **Supplementary Table 8** and meta-analytic estimates in **Supplementary Table 9**.

### Biological annotation

#### Enrichment of tissue-specific gene expression

We used gene-sets defined in Finucane et al. 2018^92^ to test for the enrichment of genes specifically expressed in one of 53 GTEx tissues^63^, or 152 tissues captured by the Franke et al. aggregation of RNA-seq studies^64,65^. This analysis seeks to confirm the role of brain tissues in mediating *Cog* and *NonCog* influences on educational attainment. The exact analysis pipeline used is available online (https://github.com/bulik/ldsc/wiki/Cell-type-specific-analyses).

#### Enrichment of cell-type specific expression

We leveraged single cell RNA sequencing (scRNA-seq) data of cells sampled from the mouse nervous system^68^ to identify cell-type specific RNA expression. Zeisel et al.^68^ sequenced cells obtained from 19 regions in the contiguous anatomical regions in the peripheral sensory, enteric, and sympathetic nervous system. After initial QC, Zeisel et al. retained 492,949 cells, which were sampled down to 160,796 high quality cells. These cells were further grouped into clusters representing 265 broad cell-types. We analyzed the dataset published by Zeisel et al. containing mean transcript counts for all genes with count >1 for each of the 265 clusters (**Supplementary Table 11**). We restricted analysis to genes with expression levels above the 25^th^ percentile. For each gene in each cell-type, we computed the cell-type specific proportion of reads for the gene (normalizing the expression within cell-type). We then computed the proportion of proportions over the 265 cell-types (computing the specificity of the gene to a specific cell-type). We ranked the 12,119 genes retained in terms of specificity to each cell-type and then retained the 10% of genes most specific to a cell-type as the “cell-type specific” gene-set. We then tested whether any of the 265 cell-type specific gene-sets were enriched in the *Cog* or *NonCog* GWAS. This analysis sought to identify specific cell-types and specific regions in the brain involved in the etiology of *Cog* and *NonCog*. We further computed the difference in enrichment for *Cog* and *NonCog* to test if any cell types were specific to either trait. For these analyses, we leveraged two widely used enrichment analysis tools: MAGMA^66^ and stratified LD score regression^67^ with the European reference panel from 1000 Genomes Project Phase 3 as SNP location and LD structure reference, Gencode release 19 as gene location reference and the human-mouse homology reference from MGI (http://www.informatics.jax.org/downloads/reports/HOM_MouseHumanSequence.rpt).

#### MAGMA

We used MAGMA (v1.07b^66^), a program for gene-set analysis based on GWAS summary statistics. We computed gene-level association statistics using a window of 10kb around the gene for both *Cog* and *NonCog*. We then used MAGMA to run a competitive gene-set analysis, using the gene p-values and gene correlation matrix (reflecting LD structure) produced in the gene-level analysis. The competitive gene-set analysis tests whether the genes within the cell-type-specific gene-set described above are more strongly associated with *Cog*/*NonCog* than other genes.

#### Stratified LDscore regression

We used LD-score regression to compute LD scores for the SNPs in each of our “cell-type specific” gene-sets. Parallel to MAGMA analysis, we added a 10kb window around each gene. We ran partitioned LD-score regression to compute the contribution of each gene-set to the heritability of *Cog* and *NonCog*. To guard against inflation, we use LD score best practices, and include the LD score baseline model (baselineL2.v2.2) in the analysis. We judged the statistical significance of the enrichment based on the p-value associated with the tau coefficient.

#### Difference in enrichment between *Cog* and *NonCog*

To compute differences in enrichment we compute a standardized difference between the per-annotation enrichment for *Cog* and *NonCog* as:

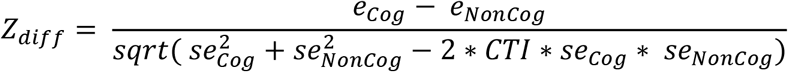

Where *e*_*Cog*_ is the enrichment of a particular gene-set for *Cog*, *e*_*NonCog*_ is the enrichment for the same gene-set for *NonCog*, *se*_*Cog*_ is the standard error of the enrichment for *Cog*, *se*_*NonCog*_ is the standard error of the enrichment for *NonCog*, and CTI is the LD score cross-trait intercept, a metric of dependence between the GWASs of *Cog* and *NonCog*.

#### Enrichment of gene expression in the brain

We performed a transcriptome-wide association study (TWAS) using Gusev et al.^69^ (FUSION: http://gusevlab.org/projects/fusion/). We used pre-computed brain-gene-expression weights available on the FUSION website, generated from 452 human individuals as part of the CommonMind Consortium. We then superimposed the bivariate distribution of the results of the TWAS for *Cog* and *NonCog* over the bivariate distribution expected given the sample overlap between EA and CP (the GWAS on which our GWAS of *Cog* and *NonCog* are based, see **Supplementary Note 2**).

### Brain modalities

#### Brain volumes

We conducted genetic correlation analysis of brain volumes using GWAS results published by Zhao et al.^72^. Zhao et al. performed GWAS of total brain volume and 100 regional brain volumes, including 99 gray matter volumes and total white matter volume (**Supplementary Table 15**). Analyses included covariate adjustment for sex, age, their square interaction and 20 principle components. Analyses of regional brain volumes additionally included covariate adjustment for total brain volume. GWAS summary statistics for these 101 brain volumes were obtained from https://med.sites.unc.edu/bigs2/data/gwas-summary-statistics/. Summary statistics were filtered and pre-processed using Genomic SEM’s “munge” function, retaining all HapMap3 SNPs with allele frequency >.01 outside the MHC region. We used Genomic-SEM to compute the genetic correlations between *Cog, NonCog* and brain volumes. Analyses of regional volumes controlled for total brain volume. For each volume, we tested if correlations differed between *Cog* and *NonCog.* Specifically, we used a chi-squared test to evaluate the null hypothesis that the two genetic correlations were equal. We used FDR adjustment to correct for multiple testing. The FDR adjustment is applied to the results for all gray matter volumes for *Cog* and *NonCog* separately.

#### White matter structures

We conducted genetic-correlation analysis of white-matter structures using GWAS results published by Zhao et al.^73^. Zhao et al. performed GWAS of diffusion tensor imaging (DTI) measures of the integrity of white-matter tracts. DTI parameters were derived for fractional anisotropy (FA), mean diffusivity (MD), axial diffusivity (AD), radial diffusivity (RD), and mode of anisotropy (MO). Each of these parameters were measured for 22 white matter tracts of interests (**Supplementary Table 16**) resulting in 110 GWAS. GWAS summary statistics for these 110 GWAS were obtained from https://med.sites.unc.edu/bigs2/data/gwas-summary-statistics/. Summary statistics were filtered and processed using Genomic SEM’s “munge” function; retaining all HapMap3 SNPs with allele frequency >.01 outside the MHC region. For each white matter structure, we tested if genetic correlations differed between *Cog* and *NonCog.* Specifically, we used a chi-squared test to evaluate the null hypothesis that the two genetic correlations were equal. We used FDR adjustment to correct for multiple testing. As these different diffusion parameters are statistically and logically interdependent, having been derived from the same tensor, FDR adjustment was applied to the results for each type of white matter diffusion parameter separately. FDR correction was applied separately for *Cog* and *NonCog*.

## Data and Resources

An FAQ on why, how and what we studied: https://medium.com/@kph3k/investigating-the-genetic-architecture-of-non-cognitive-skills-using-gwas-by-subtraction-b8743773ce44 GWAS summary data for Cog & NonCog: https://www.dropbox.com/s/cvzcedsfhbznv36/GWAS_sumstats_Cog_NonCog_Demange_et_al.zip?dl=0 A tutorial on how to perform GWAS-by-subtraction: http://rpubs.com/MichelNivard/565885

## Acknowledgements

This study was developed with support from the Jacobs Foundation at a meeting organized by DWB and KPH with support from ETD and CM and attended by PB, BWD, JW, and others. We gratefully acknowledge contributions to the meeting from Katrin Mannik and Felix Tropf, and the Jacobs Foundation Fellowship team who made the meeting possible. DWB, KPH, MGN, ETD, CM are fellows of the Foundation. JW is a Jacobs Foundation Young Scholar. We would like to thank the research participants and employees of 23andMe for making this work possible. The study also used data from the Netherlands Twin Register (NTR), the Texas Twin Study, the National Longitudinal Study of Adolescent to Adult Health (Add Health), the Dunedin Longitudinal Study, the E-Risk Study, and the Wisconsin Longitudinal Study (WLS). NTR is supported by: ‘Twin-family database for behavior genetics and genomics studies’ (NWO 480-04-004), Longitudinal data collection from teachers of Dutch twins and their siblings (NWO-481-08-011); Twin-family study of individual differences in school achievement (NWO 056-32-010) and Gravitation program of the Dutch Ministry of Education, Culture and Science and the Netherlands Organization for Scientific Research (NWO 0240-001-003); NWO Groot (480-15-001/674): Netherlands Twin Registry Repository: researching the interplay between genome and environment; NWO-Spi-56-464-14192 Biobanking and Biomolecular Resources Research Infrastructure (BBMRI – NL, 184.021.007 and 184.033.111); European Research Council (ERC-230374); the Avera Institute for Human Genetics, Sioux Falls, South Dakota (USA) and the National Institutes of Health (NIH, R01D0042157-01A); the NIMH Grand Opportunity grants (1RC2MH089951-01 and 1RC2 MH089995-01). The Texas Twin Project is supported by NICHD grants R01HD083613 and R01HD092548. Add Health is supported by Eunice Kennedy Shriver National Institute of Child Health and Human Development grant P01HD31921, and GWAS grants R01HD073342 and R01HD060726, with cooperative funding from 23 other federal agencies and foundations. The Dunedin Multidisciplinary Health and Development Study is supported by the NZ HRC, NZ MBIE, National Institute on Aging grant R01AG032282, and UK Medical Research Council grant MR/P005918/1. The E-Risk Study is supported by the UK Medical Research Council grant G1002190 and Eunice Kennedy Shriver National Institute of Child Health and Human Development grant R01HD077482. The Wisconsin Longitudinal Study is supported by National Institute on Aging grants R01AG041868 and P30AG017266.

This research received additional support from the National Institute on Aging grant R24AG045061. This research benefited from GWAS results made publicly available by the SSGAC. Some of the work used a high-performance computing facility partially supported by grant 2016-IDG-1013 from the North Carolina Biotechnology Center.

PAD is supported by the grant 531003014 from The Netherlands Organisation for Health Research and Development (ZonMW). PB is supported by the NORFACE-DIAL grant number 462-16-100.ETD is supported by NIH grants R01AG054628 and R01HD083613. The Population Research Center at the University of Texas at Austin is supported by NIH grant P2CHD042849. AA is supported by the Foundation Volksbond Rotterdam. BD is supported by award # 96-17-04 from the Russell Sage Foundation and the Ford Foundation. DIB is supported by the Royal Netherlands Academy of Science (KNAW) Professor Award (PAH/6635). HFI was supported by the “Aggression in Children: Unraveling gene-environment interplay to inform Treatment and InterventiON strategies” project (ACTION). ACTION received funding from the European Union Seventh Framework Program (FP7/2007-2013) under grant agreement no 602768. EvB is supported by NWO VENI grant 451-15-017. KPH and ETD are Faculty Research Associates of the Population Research Center at the University of Texas at Austin, which is supported by grant, 5-R24-HD042849, from the Eunice Kennedy Shriver National Institute of Child Health and Human Development (NICHD). MGN is supported by ZonMW grants 849200011 and 531003014 from The Netherlands Organisation for Health Research and Development, a VENI grant awarded by NWO (VI.Veni.191G.030).

## References

1. Moffitt, T. E. et al. A gradient of childhood self-control predicts health, wealth, and public safety. Proc. Natl. Acad. Sci. 108, 2693–2698 (2011).

2. von Stumm, S., Hell, B. & Chamorro-Premuzic, T. The Hungry Mind: Intellectual Curiosity Is the Third Pillar of Academic Performance. Perspect. Psychol. Sci. 6, 574–588 (2011).

3. Tucker-Drob, E. M., Briley, D. A., Engelhardt, L. E., Mann, F. D. & Harden, K. P. Genetically-mediated associations between measures of childhood character and academic achievement. J. Pers. Soc. Psychol. 111, 790–815 (2016).

4. Heckman, J. J., Stixrud, J. & Urzua, S. The Effects of Cognitive and Noncognitive Abilities on Labor Market Outcomes and Social Behavior. J. Labor Econ. 24, 411–482 (2006).

5. Heckman, J. J., Moon, S. H., Pinto, R., Savelyev, P. A. & Yavitz, A. The rate of return to the HighScope Perry Preschool Program. J. Public Econ. 94, 114–128 (2010).

6. Conti, G., Heckman, J. J. & Pinto, R. The Effects of Two Influential Early Childhood Interventions on Health and Healthy Behaviour. Econ. J. 126, F28–F65 (2016).

7. Gutman, L. M. & Schoon, I. The impact of non-cognitive skills on outcomes for young people. Educ. Endow. Found. 59, 2019 (2013).

8. Garcia, E. The Need to Address Noncognitive Skills in the Education Policy Agenda. https://www.epi.org/publication/the-need-to-address-noncognitive-skills-in-the-education-policy-agenda/ (2014).

9. Kautz, T., Heckman, J. J., Diris, R., Ter Weel, B. & Borghans, L. Fostering and Measuring Skills: Improving Cognitive and Non-Cognitive Skills to Promote Lifetime Success.” OECD Education Working Papers, No. 110, OECD Publishing, Paris. (2014).

10. Heckman, J. J. Skill Formation and the Economics of Investing in Disadvantaged Children. LIFE CYCLES 312, 4 (2006).

11. Heckman, J. J. & Kautz, T. Hard evidence on soft skills. Labour Econ. 19, 451–464 (2012).

12. Rimfeld, K., Kovas, Y., Dale, P. S. & Plomin, R. True grit and genetics: Predicting academic achievement from personality. J. Pers. Soc. Psychol. 111, 780–789 (2016).

13. Richardson, M., Abraham, C. & Bond, R. Psychological correlates of university students’ academic performance: A systematic review and meta-analysis. Psychol. Bull. 138, 353–387 (2012).

14. Smithers, L. G. et al. A systematic review and meta-analysis of effects of early life non-cognitive skills on academic, psychosocial, cognitive and health outcomes. Nat. Hum. Behav. 2, 867–880 (2018).

15. Kovas, Y. et al. Why children differ in motivation to learn: Insights from over 13,000 twins from 6 countries. Personal. Individ. Differ. 80, 51–63 (2015).

16. Loehlin, J. C. Genes and environment in personality development. (Sage Publications, 1992).

17. Tucker-Drob, E. M. & Harden, K. P. Learning motivation mediates gene-by-socioeconomic status interaction on mathematics achievement in early childhood. Learn. Individ. Differ. 22, 37–45 (2012).

18. Malanchini, M., Engelhardt, L. E., Grotzinger, A. D., Harden, K. P. & Tucker-Drob, E. M. “Same but different”: Associations between multiple aspects of self-regulation, cognition, and academic abilities. J. Pers. Soc. Psychol. 117, 1164–1188 (2019).

19. Morris, T. T., Smith, G. D., van Den Berg, G. & Davies, N. M. Investigating the longitudinal consistency and genetic architecture of non-cognitive skills, and their relation to educational attainment. http://biorxiv.org/lookup/doi/10.1101/470682 (2018) doi:10.1101/470682.

20. Bowles, S. & Gintis, H. Schooling In Capitalist America: Educational Reform And The Contradictions Of Economic Life. (Basic Books, 1977).

21. Heckman, J. J. & Rubinstein, Y. The Importance of Noncognitive Skills: Lessons from the GED Testing Program. Am. Econ. Rev. 91, 145–149 (2001).

22. Grotzinger, A. D. et al. Genomic structural equation modelling provides insights into the multivariate genetic architecture of complex traits. Nat. Hum. Behav. 3, 513 (2019).

23. Lee, J. J. et al. Gene discovery and polygenic prediction from a genome-wide association study of educational attainment in 1.1 million individuals. Nat. Genet. 50, 1112–1121 (2018).

24. Ligthart, L. et al. The Netherlands Twin Register: Longitudinal Research Based on Twin and Twin-Family Designs. Twin Res. Hum. Genet. 1–14 (2019) doi:10.1017/thg.2019.93.

25. Harden, K. P., Tucker-Drob, E. M. & Tackett, J. L. The Texas Twin Project. Twin Res. Hum. Genet. 16, 385–390 (2013).

26. Harris, K. M. et al. Cohort Profile: The National Longitudinal Study of Adolescent to Adult Health (Add Health). Int. J. Epidemiol. 48, 1415–1415k (2019).

27. Herd, P., Carr, D. & Roan, C. Cohort profile: Wisconsin longitudinal study (WLS). Int. J. Epidemiol. 43, 34–41 (2014).

28. Poulton, R., Moffitt, T. E. & Silva, P. A. The Dunedin Multidisciplinary Health and Development Study: overview of the first 40 years, with an eye to the future. Soc. Psychiatry Psychiatr. Epidemiol. 50, 679–693 (2015).

29. Moffitt, T. E. & E-risk Team. Teen-aged mothers in contemporary Britain. J. Child Psychol. Psychiatry 43, 727–742 (2002).

30. Benyamin, B. et al. Childhood intelligence is heritable, highly polygenic and associated with FNBP1L. Mol. Psychiatry 19, 253–258 (2014).

31. Chetty, R. et al. The Association Between Income and Life Expectancy in the United States, 2001-2014. JAMA 315, 1750 (2016).

32. Case, A. & Deaton, A. Mortality and Morbidity in the 21st Century. Brook. Pap. Econ. Act. 2017, 397–476 (2017).

33. Hill, W. D. et al. Molecular Genetic Contributions to Social Deprivation and Household Income in UK Biobank. Curr. Biol. 26, 3083–3089 (2016).

34. Pilling, L. C. et al. Human longevity is influenced by many genetic variants: evidence from 75,000 UK Biobank participants. 8, 14 (2016).

35. Almlund, M., Duckworth, A. L., Heckman, J. & Kautz, T. Personality Psychology and Economics. in Handbook of the Economics of Education vol. 4 1–181 (Elsevier, 2011).

36. Borghans, L., Duckworth, A. L. & Heckman, J. J. The Economics and Psychology of Personality Traits. 88.

37. Rabin, M. A perspective on psychology and economics. Eur. Econ. Rev. 29 (2002).

38. Becker, A., Deckers, T., Dohmen, T., Falk, A. & Kosse, F. The Relationship Between Economic Preferences and Psychological Personality Measures. Annu. Rev. Econ. 4, 453–478 (2012).

39. Linnér, R. K. et al. Genome-wide association analyses of risk tolerance and risky behaviors in over 1 million individuals identify hundreds of loci and shared genetic influences. Nat. Genet. 51, 245–257 (2019).

40. Sanchez-Roige, S. et al. Genome-wide association study of delay discounting in 23,217 adult research participants of European ancestry. Nat. Neurosci. 21, 16–18 (2018).

41. Yengo, L. et al. Meta-analysis of genome-wide association studies for height and body mass index in ~700000 individuals of European ancestry. Hum. Mol. Genet. 27, 3641–3649 (2018).

42. Furberg, H., Kim, Y., Dackor, J. & Boerwinckle, E. Genome-wide meta-analyses identify multiple loci associated with smoking behavior. Nat. Genet. 42, 441–447 (2010).

43. Walters, R. K. et al. Trans-ancestral GWAS of alcohol dependence reveals common genetic underpinnings with psychiatric disorders. 34.

44. Schumann, G. et al. KLB is associated with alcohol drinking, and its gene product β-Klotho is necessary for FGF21 regulation of alcohol preference. Proc. Natl. Acad. Sci. 113, 14372–14377 (2016).

45. Pasman, J. A. et al. GWAS of lifetime cannabis use reveals new risk loci, genetic overlap with psychiatric traits, and a causal effect of schizophrenia liability. Nat. Neurosci. 21, 1161–1170 (2018).

46. Linnér, R. K., Dick, D. & Koellinger, P. D. the Externalizing Consortium – Multivariate analyses of large-scale GWAS to identify genetic factors in externalizing behaviors and disorders. (2018) doi:None.

47. Barban, N. et al. Genome-wide analysis identifies 12 loci influencing human reproductive behavior. Nat. Genet. 48, 1462–1472 (2016).

48. Lo, M.-T. et al. Genome-wide analyses for personality traits identify six genomic loci and show correlations with psychiatric disorders. Nat. Genet. 49, 152–156 (2017).

49. John, O. P., Naumann, L. P. & Soto, C. J. Paradigm shift to the integrative Big Five Trait taxonomy. Handb. Personal. Theory Res. 114–158 (2008) doi:10.1016/S0191-8869(97)81000-8.

50. de Moor, M. H. M. et al. Meta-analysis of genome-wide association studies for personality. Mol. Psychiatry 17, 337–349 (2012).

51. Caspi, A., Roberts, B. W. & Shiner, R. L. Personality development: stability and change. Annu. Rev. Psychol. 56, 453–484 (2005).

52. Kessler, R. C. et al. Social consequences of psychiatric disorders, I: educational attainment. Am. J. Psychiatry 1026–1032 (1995).

53. Breslau, J., Lane, M., Sampson, N. & Kessler, R. C. Mental Disorders and Subsequent Educational Attainment in a US National Sample. J. Psychiatr. Res. 42, 708–716 (2008).

54. Power, R. A. et al. Polygenic risk scores for schizophrenia and bipolar disorder predict creativity. Nat. Neurosci. 18, 953–955 (2015).

55. Bansal, V. et al. Genome-wide association study results for educational attainment aid in identifying genetic heterogeneity of schizophrenia. Nat. Commun. Lond. 9, 1–12 (2018).

56. Wray, N. R. et al. Genome-wide association analyses identify 44 risk variants and refine the genetic architecture of major depression. Nat. Genet. 50, 668–681 (2018).

57. Schizophrenia Working Group of the Psychiatric Genomics Consortium et al. Biological insights from 108 schizophrenia-associated genetic loci. Nature 511, 421–427 (2014).

58. Ruderfer, D. M. et al. Genomic Dissection of Bipolar Disorder and Schizophrenia, Including 28 Subphenotypes. Cell 173, 1705–1715.e16 (2018).

59. Jansen, P. R. et al. Genome-wide analysis of insomnia in 1,331,010 individuals identifies new risk loci and functional pathways. Nat. Genet. 51, 394–403 (2019).

60. Duncan, L. et al. Significant Locus and Metabolic Genetic Correlations Revealed in Genome-Wide Association Study of Anorexia Nervosa. Am. J. Psychiatry 174, 850–858 (2017).

61. International Obsessive Compulsive Disorder Foundation Genetics Collaborative (IOCDF-GC) and OCD Collaborative Genetics Association Studies (OCGAS) et al. Revealing the complex genetic architecture of obsessive–compulsive disorder using meta-analysis. Mol. Psychiatry 23, 1181–1188 (2018).

62. Grove, J. et al. Common risk variants identified in autism spectrum disorder. bioRxiv 224774 (2017) doi:10.1101/224774.

63. The GTEx Consortium et al. The Genotype-Tissue Expression (GTEx) pilot analysis: Multitissue gene regulation in humans. Science 348, 648–660 (2015).

64. Pers, T. H. et al. Biological interpretation of genome-wide association studies using predicted gene functions. Nat. Commun. 6, 5890 (2015).

65. Fehrmann, R. S. N. et al. Gene expression analysis identifies global gene dosage sensitivity in cancer. Nat. Genet. 47, 115–125 (2015).

66. de Leeuw, C. A., Mooij, J. M., Heskes, T. & Posthuma, D. MAGMA: Generalized Gene-Set Analysis of GWAS Data. PLoS Comput. Biol. 11, 1–19 (2015).

67. Finucane, H. K. et al. Partitioning heritability by functional annotation using genome-wide association summary statistics. Nat. Genet. 47, 1228–1235 (2015).

68. Zeisel, A. et al. Molecular Architecture of the Mouse Nervous System. Cell 174, 999–1014.e22 (2018).

69. Gusev, A. et al. Integrative approaches for large-scale transcriptome-wide association studies. Nat. Genet. 48, 245–252 (2016).

70. Nave, G., Jung, W. H., Karlsson Linnér, R., Kable, J. W. & Koellinger, P. D. Are Bigger Brains Smarter? Evidence From a Large-Scale Preregistered Study. Psychol. Sci. 30, 43–54 (2019).

71. Elliott, M. L. et al. A Polygenic Score for Higher Educational Attainment is Associated with Larger Brains. Cereb. Cortex 29, 3496–3504 (2019).

72. Zhao, B. et al. GWAS of 19,629 individuals identifies novel genetic variants for regional brain volumes and refines their genetic co-architecture with cognitive and mental health traits. bioRxiv 586339 (2019) doi:10.1101/586339.

73. Zhao, B. et al. Large-scale GWAS reveals genetic architecture of brain white matter microstructure and genetic overlap with cognitive and mental health traits (n=17,706). bioRxiv 288555 (2019) doi:10.1101/288555.

74. Haushofer, J. & Fehr, E. On the psychology of poverty. Science 344, 862–867 (2014).

75. Briley, D. A., Domiteaux, M. & Tucker-Drob, E. M. Achievement-relevant personality: Relations with the Big Five and validation of an efficient instrument. Learn. Individ. Differ. 32, 26–39 (2014).

76. Smoller, J. W. et al. Psychiatric genetics and the structure of psychopathology. Mol. Psychiatry 24, 409–420 (2019).

77. Plomin, R., Haworth, C. M. A. & Davis, O. S. P. Common disorders are quantitative traits. Nat. Rev. Genet. 10, 872–878 (2009).

78. Meehl, P. E. Schizotaxia, schizotypy, schizophrenia. Am. Psychol. 17, 827–838 (1962).

79. Jencks, C. Inequality: A reassessment of the effect of family and schooling in America. (Basic Books, 1972).

80. von Stumm, S. & Ackerman, P. L. Investment and intellect: A review and meta-analysis. Psychol. Bull. 139, 841–869 (2013).

81. Tucker-Drob, E. M. & Harden, K. P. A Behavioral Genetic Perspective on Non-Cognitive Factors and Academic Achievement. in Genetics, Ethics and Education (eds. Grigorenko, E. L., Tan, M., Latham, S. R. & Bouregy, S.) 134–158 (Cambridge University Press, 2017). doi:10.1017/9781316340301.007.

82. Tucker-Drob, E. M. Motivational factors as mechanisms of gene-environment transactions in cognitive development and academic achievement. in Handbook of competence and motivation: Theory and application, 2nd ed. 471–486 (The Guilford Press, 2017).

83. Tucker-Drob, E. M. & Harden, K. P. Intellectual Interest Mediates Gene × Socioeconomic Status Interaction on Adolescent Academic Achievement: Intellectual Interest and G×E. Child Dev. no-no (2012) doi:10.1111/j.1467-8624.2011.01721.x.

84. Malanchini, M. et al. Reading self-perceived ability, enjoyment and achievement: A genetically informative study of their reciprocal links over time. Dev. Psychol. 53, 698–712 (2017).

85. Tropf, F. C. et al. Hidden heritability due to heterogeneity across seven populations. Nat. Hum. Behav. 1, 757–765 (2017).

86. Duncan, L. et al. Analysis of polygenic risk score usage and performance in diverse human populations. Nat. Commun. 10, 3328 (2019).

87. Martin, A. R. et al. Human Demographic History Impacts Genetic Risk Prediction across Diverse Populations. Am. J. Hum. Genet. 100, 635–649 (2017).

88. Willer, C. J., Li, Y. & Abecasis, G. R. METAL: Fast and efficient meta-analysis of genomewide association scans. Bioinformatics 26, 2190–2191 (2010).

89. Willemsen, G. et al. The Adult Netherlands Twin Register: Twenty-Five Years of Survey and Biological Data Collection. Twin Res. Hum. Genet. 16, 271–281 (2013).

90. Highland, H. M., Avery, C. L., Duan, Q., Li, Y. & Harris, K. M. Quality control analysis of Add Health GWAS data. https://www.cpc.unc.edu/projects/addhealth/documentation/guides/AH_GWAS_QC.pdf (2018).

91. Vilhjálmsson, B. J. et al. Modeling Linkage Disequilibrium Increases Accuracy of Polygenic Risk Scores. Am. J. Hum. Genet. 97, 576–592 (2015).

92. Finucane, H. K. et al. Heritability enrichment of specifically expressed genes identifies disease-relevant tissues and cell types. Nat. Genet. 50, 621–629 (2018).

