## Supplementary notes for "Investigating the Genetic Architecture of Non-Cognitive Skills Using GWAS-by-Subtraction"

### Supplementary Note 1. Comparison Cog GWAS and original Cognitive Performance GWAS

We defined our genomic structural equation model such that the latent variable *Cog* is equivalent to the Cognitive Performance variable. We therefore expected our *Cog* GWAS to parallel the original Cognitive Performance GWAS[1]. We confirm that both GWAS are highly correlated with a LDSC genetic correlation of 1 (*p*=.00). Both have a similar SNP heritability: *Cog* SNP-$h^2$= .1921 (*SE*=.0061) and CP SNP-$h^2$=.192 (*SE*=.0062). We found 282 significantly associated hits with *Cog* (**Supplementary Figure 1**), while Lee et al. report only 225 significant hits. However, Lee et al. used a different pruning method: they did not use a distance threshold. Re-analysing their summary statistics with our pruning method (radius of 250kb and LD threshold of $r^2 < 0.1$), we found 294 independent SNPs associated with CP. The discrepancy between the 282 hits associated with *Cog* and 294 with CP is due to the difference of SNPs present in both GWAS (7 311 269 for *Cog* and 10 098 325 for CP).

### Supplementary Note 2. Cell-type enrichment with Stratified LDSC regression

Additionally to MAGMA[2] analysis, we tested for cell-type specific gene-sets enrichments with stratified LDSC regression[3]. Enrichment in MAGMA tests the enrichment of a particular gene-set relative to all genes. In contrast, enrichment in LD score regression implies enrichment over and above the signal present in any of the “baseline” annotation, which include genes, promoters, histone marks, and other salient features of the genome. Using LDSC partitioned heritability, we found similar results as with MAGMA. The Spearman rank correlation between –log(*p*) of MAGMA estimate and LDSC Enrichment[4] was 0.89 for *NonCog* and 0.83 for *Cog*, but fewer cell-types are significantly enriched (**Supplementary Figure 6** and **Table 13**). For the *NonCog* factor, only one depletion was found for vascular cell (VECC). As with MAGMA, the correlation between the LD score regression *Z*-statistics for *Cog* and *NonCog* was substantial (*r*=.62), and there was no significant difference in cell-type specific enrichment between the two factors.

### Supplementary Note 3. Transcriptome-wide association study

It is somewhat remarkable that while *Cog* and *NonCog* are genetically uncorrelated by design, the heritable signal for either is enriched in the same pathways at the cell-type level. This points to enrichment in the same pathways being driven by distinct sets of genes. We further investigated the differentiation of *Cog* and *NonCog* at the level of the gene. A transcriptome-wide association study[5] revealed that the set of genes associated with *Cog* and *NonCog* differ (**Supplementary Table 14**). The correlation between the Z-statistics for the association with *NonCog* and *Cog* was negative (-0.3). This result is in line with the fact that the samples for cognitive performance and educational attainment overlap. Indeed, the LD score cross-trait intercept between *Cog* and *NonCog* was estimated at -0.67. This represents the dependence of the results between our two GWAS induced by sample overlap between the initial GWAS of education and cognitive performance. The presence of a correlation introduced by sample overlap can bias our results if we do not consider it when evaluating TWAS results. We plot the distribution of the expected dependence under the null and superimpose the true TWAS results for *Cog* and *NonCog* in **Supplementary Figure 7**. The blue ellipses correspond to 68% and 95% of the bivariate density under the null, while the red line corresponds to a density of 1-(0.05/5378), representing the gene wide significance level of this null-distribution. The result is consistent with TWAS results being a function of the sample overlap, with true signal departing from the null-distribution both in concordant and discordant directions. We observe no apparent correlation between the Z-statistics for the association between gene expression for *Cog* and *NonCog*, beyond the effects of sample overlap. We observe significant associations between genes and *NonCog* and *Cog* respectively in concordant direction, as well as discordant direction.

### Supplementary references


1. Lee, J. J. *et al.* Gene discovery and polygenic prediction from a genome-wide association study of educational attainment in 1.1 million individuals. *Nat. Genet.* **50**, 1112–1121 (2018).
2. de Leeuw, C. A., Mooij, J. M., Heskes, T. & Posthuma, D. MAGMA: Generalized Gene-Set Analysis of GWAS Data. *PLOS Comput. Biol.* **11**, e1004219 (2015).
3. Finucane, H. K. *et al.* Partitioning heritability by functional annotation using genome-wide association summary statistics. *Nat. Genet.* **47**, 1228–1235 (2015).
4. Bryois, J. *et al. Genetic Identification of Cell Types Underlying Brain Complex Traits Yields Novel Insights Into the Etiology of Parkinson's Disease*. http://biorxiv.org/lookup/doi/10.1101/528463 (2019) doi:10.1101/528463.
5. Gusev, A. *et al.* Integrative approaches for large-scale transcriptome-wide association studies. *Nat. Genet.* **48**, 245–252 (2016).
