## Supplementary figures for "Investigating the Genetic Architecture of Non-Cognitive Skills Using GWAS-by-Subtraction"

### Supplementary Figure 1. Manhattan plot of the *Cog* GWAS

Plot of the -log$_{10}$(p-value) associated with the Wald test of $\beta_{Cog}$ for all SNPs ordered by chromosome and base position. Purple triangles indicate genome-wide significant ($p < 5e10^{-8}$) and independent (within a 250Kb window and $r^2 < .1$) associations.

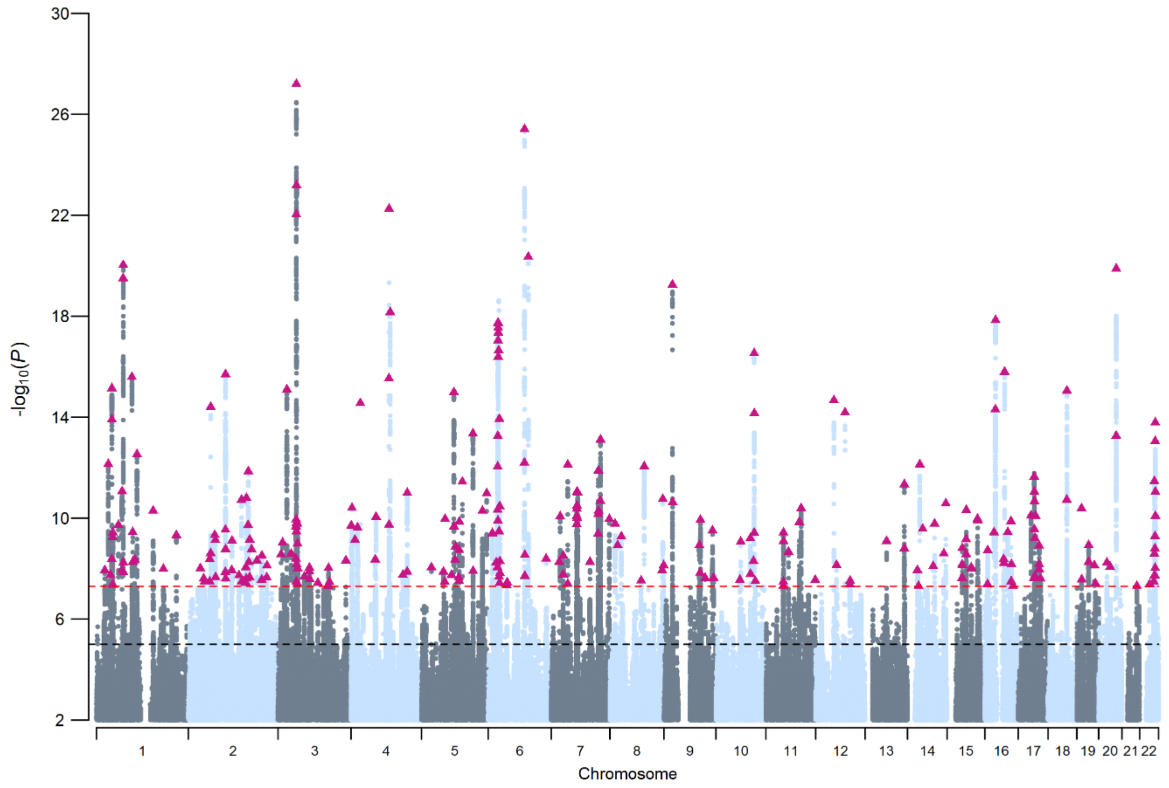

Supplementary Figure 2. PGS predictions of cognitive performance

Meta-analytic estimates of the polygenic score associations with cognitive test performance. *Cog* and *NonCog* PRS were entered simultaneously in multiple regression. Results are ordered and meta-analysed separately for fluid (top panel) and crystalized (bottom panel) intelligence tests. Width of the band represents 95% CI.

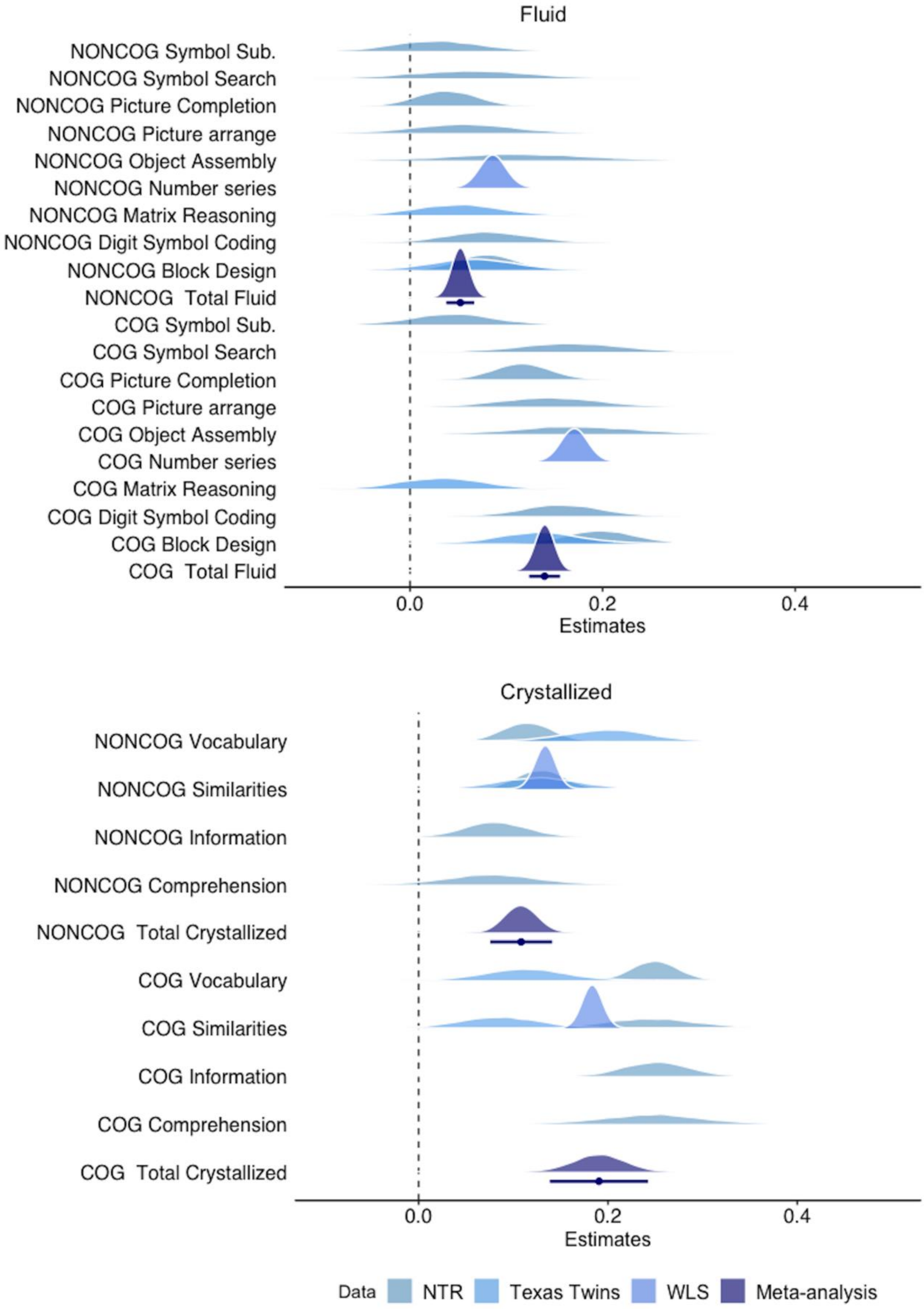

Supplementary Figure 3. PGS predictions of personality traits

Meta-analytic estimates of the polygenic score associations with Big-5 personality traits. *Cog* and *NonCog* PRS were entered simultaneously in multiple regression.

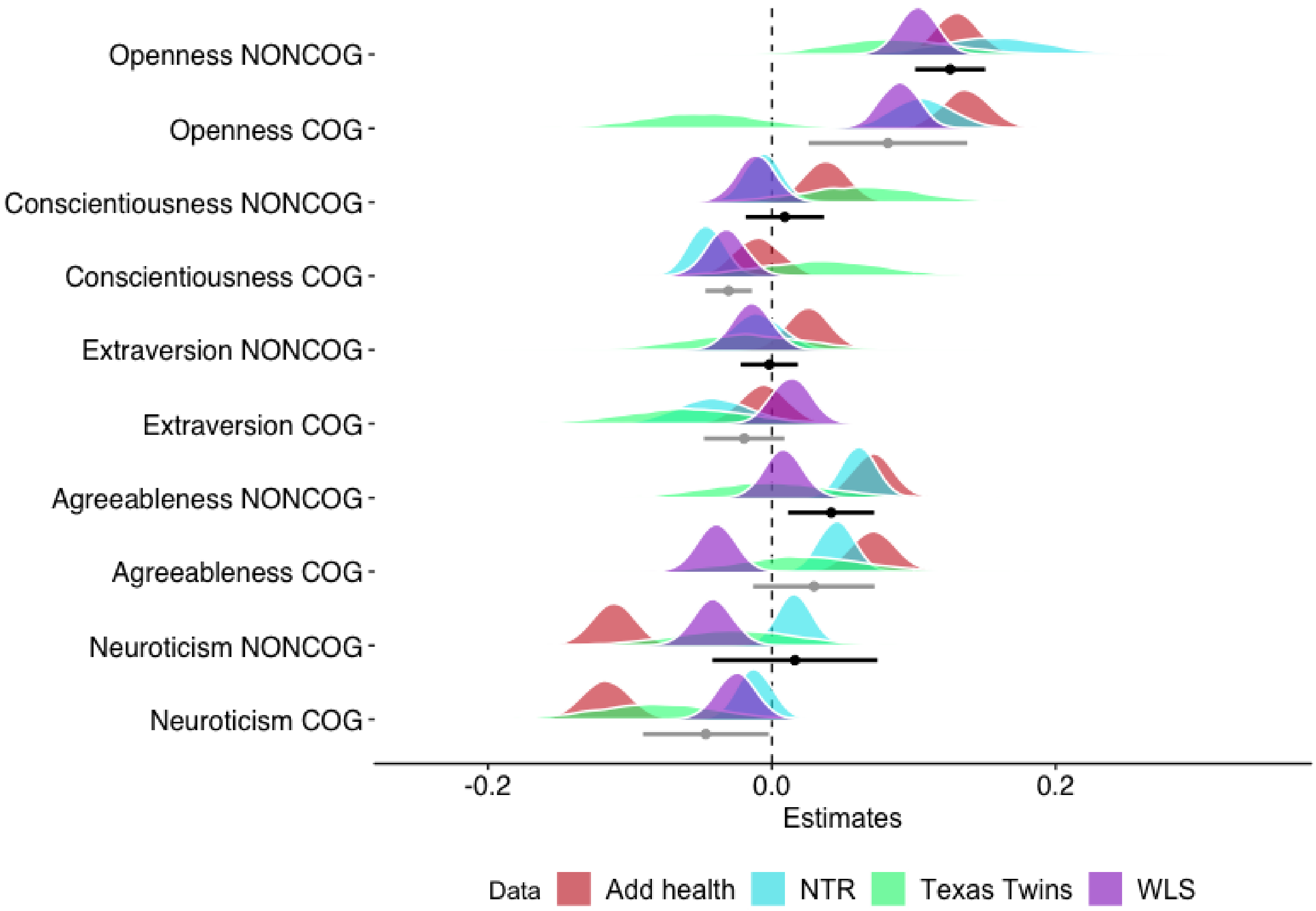

Supplementary Figure 4. Z-scores of enrichment of *Cog* and *NonCog* for 265 cell-types computed with MAGMA

Categories of the nervous system cell-types are defined following Taxonomy 2 (Supplementary Table 11). Associations significant after FDR correction are represented in orange, in red if significant after Bonferroni correction.

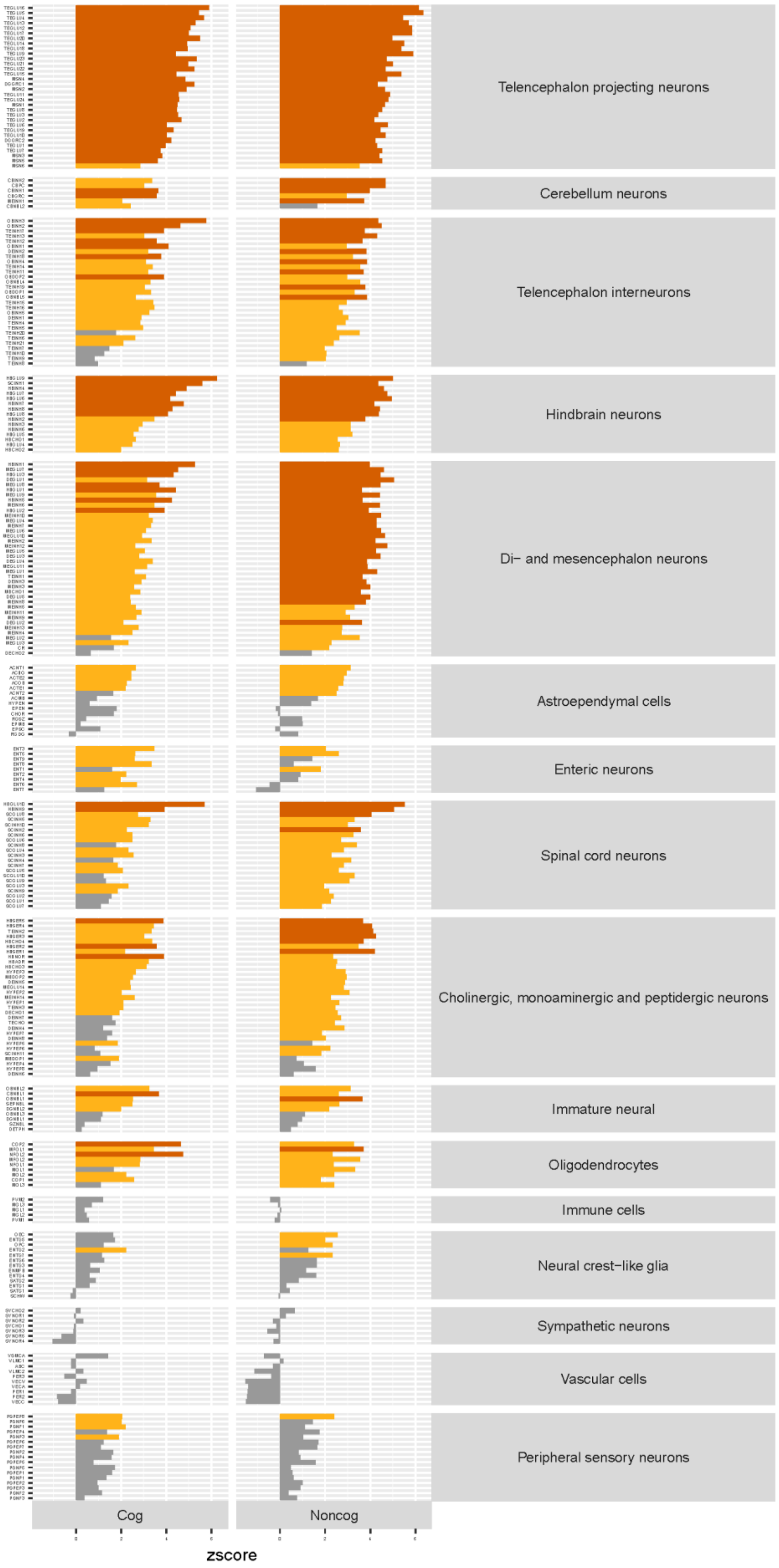

Supplementary Figure 5. Plot of MAGMA cell-types enrichment z-scores of *Cog* vs *NonCog*

Cell-types are categorized following Taxonomy 1 (Supplementary Table 11). Black lines represent significant thresholds with Bonferroni correction.

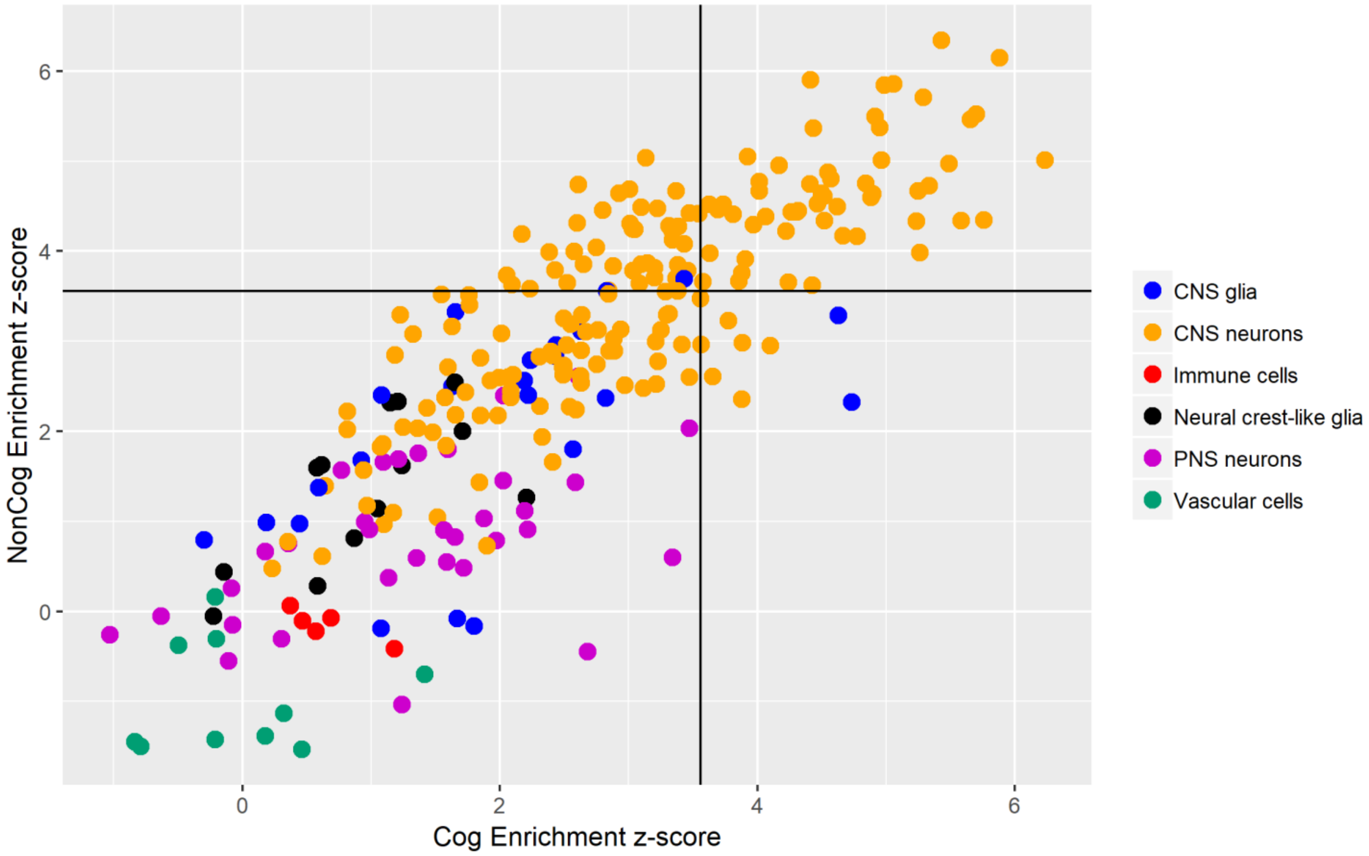

Supplementary Figure 6. Z-scores of enrichment of Cog and NonCog for 265 cell-types computed with stratified LDSC regression

Categories of the nervous system cell-types are defined following Taxonomy 2 (Supplementary Table 11). Associations significant after FDR correction are represented in orange, in red if significant after Bonferroni correction.

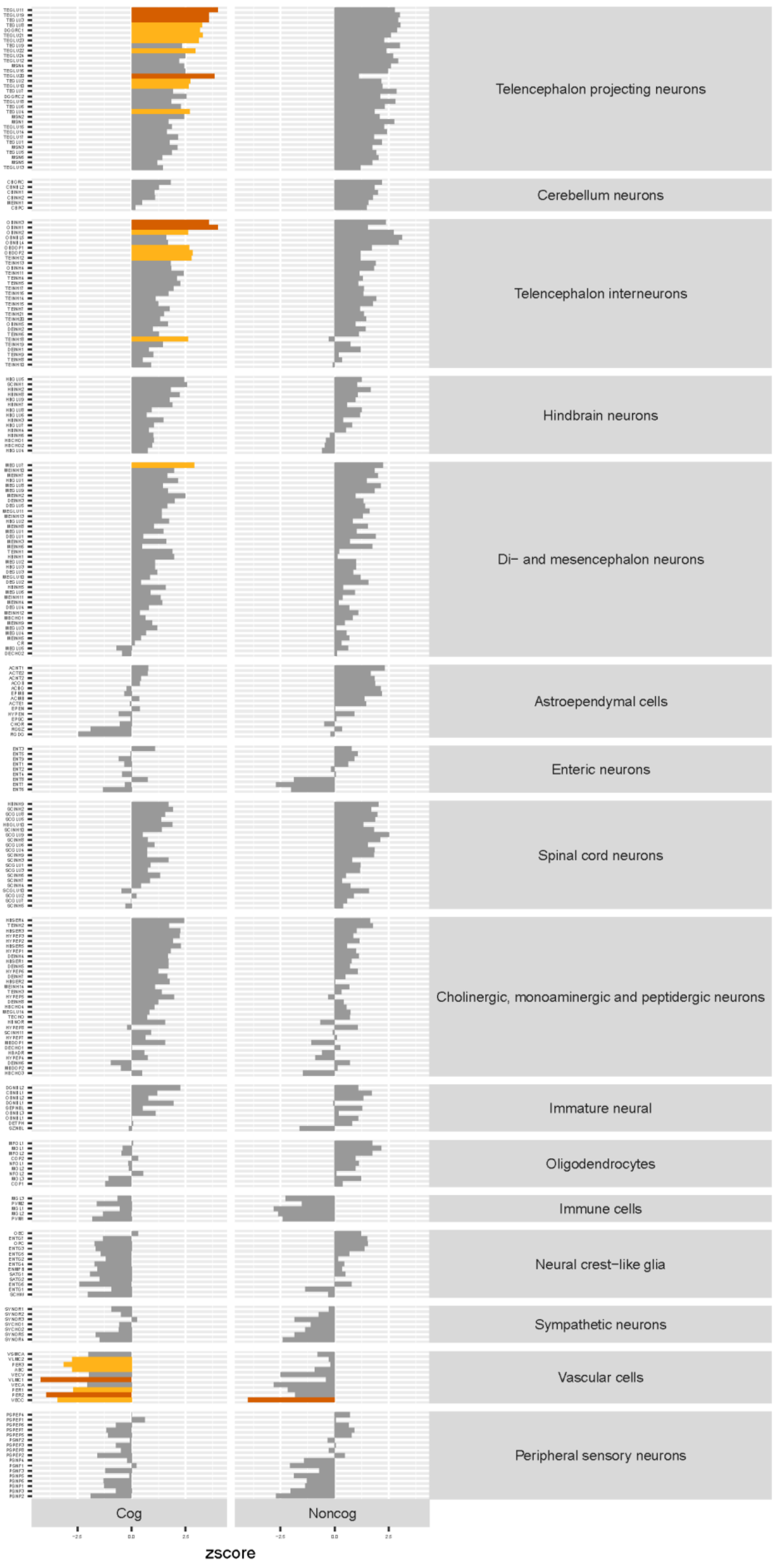

Supplementary Figure 7.  Scatterplot of TWAS Z-scores for *Cog* and *NonCog*

Scatterplot of gene test-statistics for its effects on *Cog* and *NonCog*. Blue ellipses represent the 68% and 95% CI of the bivariate null-distribution of test-statistics (as expected given the LD score cross-trait intercept for *Cog* and *NonCog* of -0.67). Red ellipse represent the gene-wide significance level of this bivariate null-distribution ($p < 0.05$/N genes in TWAS). The histograms represent the distributions of Z-scores for all genes considered in the TWAS, with overline of the standard normal distribution in blue.

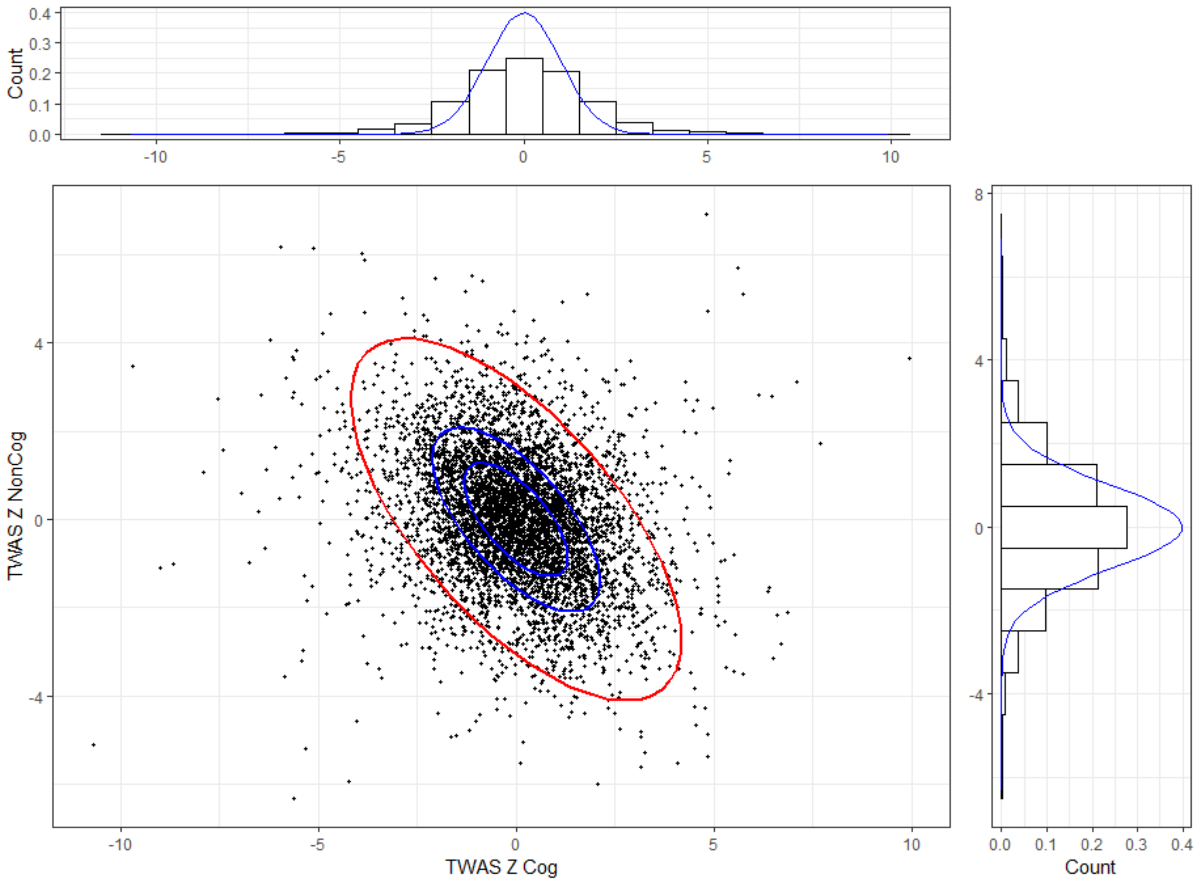

Supplementary Figure 8. Genetic correlation using Genomic SEM without SNP effects

Genomic SEM model used to directly estimate the covariance between *Cog* and *NonCog* and a third trait, without the need to first perform a *Cog* and *NonCog* GWAS. The covariance of EA and a tertiary trait explained by *NonCog* is calculated as ENC = λNonCog-EA * λNonCog-LatentTrait * λTrait. The covariance between EA and a tertiary trait explained by *Cog* is calculated as EC = λCog-EA * λCog-LatentTrait * λTrait. The total covariance between EA and a tertiary trait is calculated as Etotal = λNonCog-EA * λNonCog-LatentTrait * λTrait + λCog-EA * λCog-LatentTrait * λTrait. The percentage of the genetic covariance explained by *NonCog* was ENC/Etotal and the percentage explained by *Cog* was defined as EC/Etotal. This analysis only yields valid results for trait where ENC and EC are both positive or both negative.

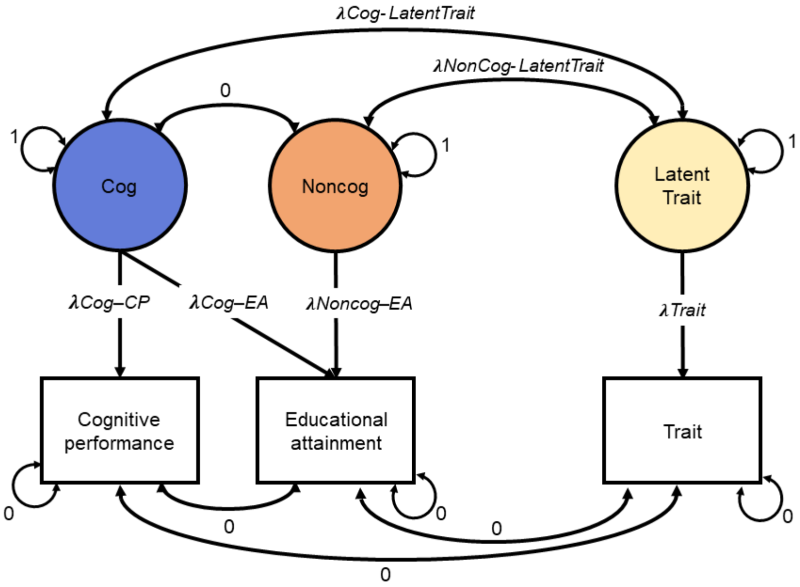
